# Perlin Noise Generation of Physiologically Realistic Patterns of Fibrosis

**DOI:** 10.1101/668848

**Authors:** David Jakes, Kevin Burrage, Christopher C. Drovandi, Pamela Burrage, Alfonso Bueno-Orovio, Rodrigo Weber dos Santos, Blanca Rodriguez, Brodie A. J. Lawson

## Abstract

Fibrosis, the pathological excess of fibroblast activity, is a significant health issue that hinders the function of many organs in the body, in some cases fatally. However, the severity of fibrosis-derived conditions depends on both the positioning of fibrotic affliction, and the microscopic patterning of fibroblast-deposited matrix proteins within afflicted regions. Variability in an individual’s manifestation of a type of fibrosis is an important factor in explaining differences in symptoms, optimum treatment and prognosis, but a need for *ex vivo* procedures and a lack of experimental control over conflating factors has meant this variability remains poorly understood. In this work, we present a computational methodology for the generation of patterns of fibrosis microstructure, demonstrating the technique using histological images of four types of cardiac fibrosis. Our generator and automated tuning method prove flexible enough to capture each of these very distinct patterns, allowing for rapid generation of new realisations for high-throughput computational studies. We also demonstrate via simulation, using the generated fibrotic patterns, the importance of micro-scale variability by showing significant differences in electrophysiological impact even within a single class of fibrosis.

## Introduction

Fibrosis, the excess of fibroblast activity resulting in a pathological accumulation of extracellular matrix proteins such as collagen [1], is a condition that significantly (in many cases, eventually fatally) compromises the function of many organs of the body, in particular the heart [2], lungs [3] and liver [4]. Owing to the different triggers for fibroblast activity, there exist many different variations of fibrosis specific to the organs they impact, often characterised by the spatial arrangement of afflicted tissue [5, 6]. Beyond these broad categories, the specific distribution of fibrosis on the micro-scale also affects its impact on organ function, particularly in the case of the heart where specific micro-anatomical features can act to create dangerous re-entrant arrhythmias [7].

Non-invasive imaging techniques remain unable to fully characterise the nature of fibrosis in the lung [8] and particularly the heart [9], resulting in a reliance upon *ex vivo* studies [10, 11]. This has motivated *in silico* investigations, seeking to explain both the creation of different fibrotic patternings [12] and in the case of cardiac fibrosis, its pro-arrhythmic consequences [13, 14, 15, 16, 17]. Although some *in silico* investigations have directly used invasively-collected data of cardiac fibrosis localisation to study the electrophysiological impacts of realistic patterns of affliction [10, 18, 19, 20], the reliance on this limited data has meant that the effects of variability in these patterns remains very poorly understood.

In this work, we address this issue by presenting a framework for the computational generation of physiologically accurate patterns of fibrosis. We work directly from experimental images of collagen distribution in four different classifications of cardiac fibrosis, constructing a quantitative measure of matching between patterns that considers their similarity in terms of key pattern features. This allows us to generate new pattern realisations that successfully capture the distinguishing features of interstitial, compact, diffuse and patchy fibrosis [5] (Figure 1).

**Figure 1:**
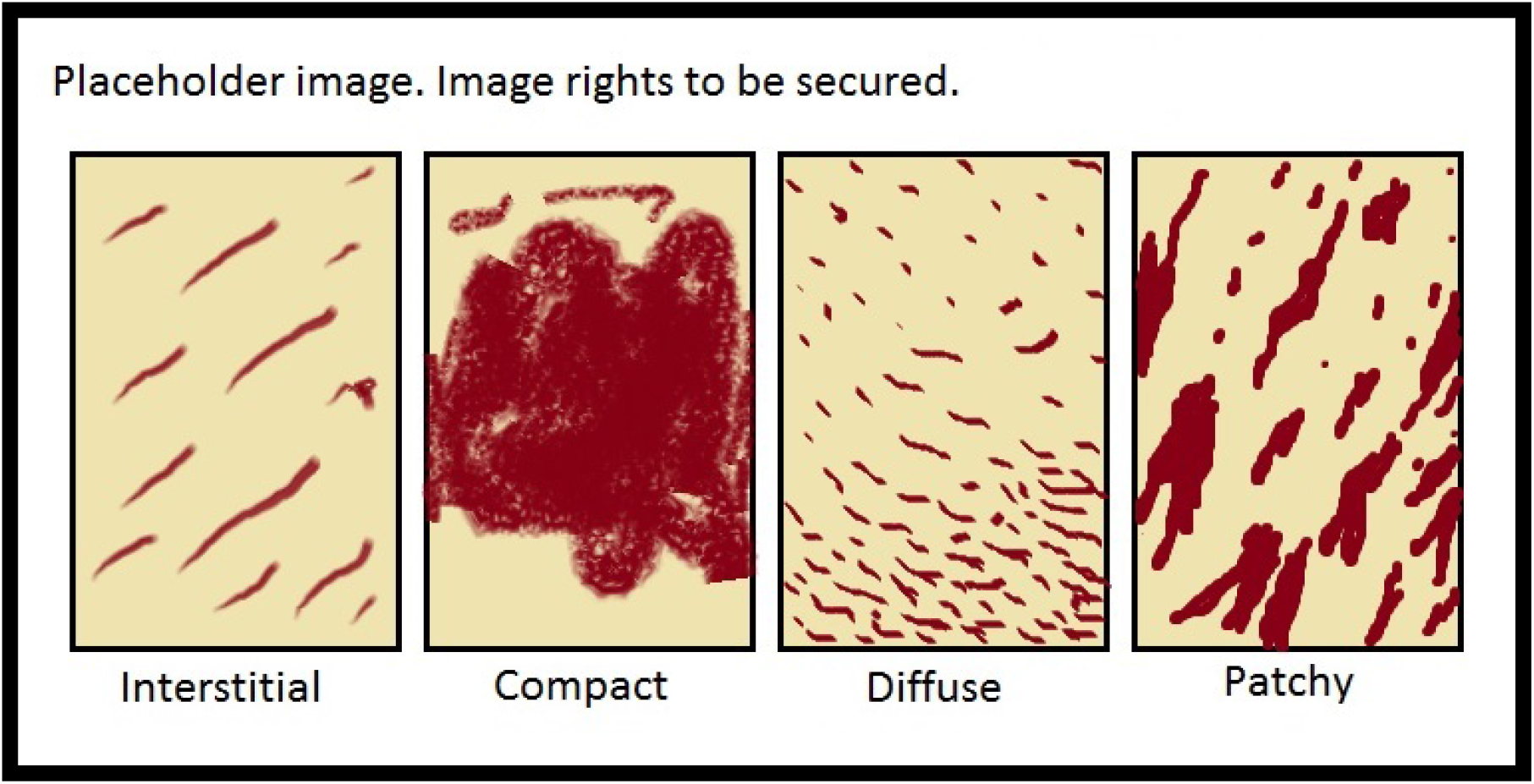
Image rights still to be secured at time of publication. To be visualised are the four types of microfibrosis as presented by de Jong *et al.* [5], converted to binary images via the thresholding process described in Methods. This image simply demonstrates the distinctive features of the four classes in a rough illustrative fashion.

Our pattern generation methodology, leveraging the Perlin noise technique for computationally-efficient generation of textures in computer graphics [21], is flexible enough to create a wide range of patterns and controlled by a set of parameters that each has a physical interpretation, allowing for intuitive tuning. Furthermore, we demonstrate how approximate Bayesian computation (ABC) can be exploited to create a tuning-free method for the automatic generation of a population of patterns matched to a provided target. Perlin noise [22] and a different type of noisefield, Gaussian random fields [23], have been used to generate patterns of fibrosis in breast tissue and the heart, respectively. However, no previous works have attempted to use such an approach to capture the intricate patterns observed on the microscopic scale, where such techniques are arguably most needed.

We demonstrate the efficacy of our generator and tuning techniques by first re-creating artificially-generated target patterns, and then the patterns evident in histological images of cardiac fibrosis. We also use our computationally-generated patterns to highlight the importance of the studies into variability that our techniques enable, demonstrating via simulation the significant differences in the impacts on cardiac excitation propagation that occur even between multiple realisations of the same class of fibrosis. We conclude by discussing further potential applications for our methodology, including its extension to macroscopic patternings on realistic anatomies.

## Results

### Perlin Noise Combinations Offer Controllable, Flexible Pattern Generation

We use a combination of Perlin noises to generate “images” with continuous-valued pixels, which are then thresholded to create binary patterns representing the presence or absence of fibrotic obstruction. A primary appeal of pattern generation by noisefield in general is that many patterns with the same features can be created simply by generating many realisations of the noisefield. Our pattern generator (described in Methods) is composed of a weighted combination of three separate fields created using Perlin noise, each with a distinct purpose and its own set of associated parameters, as summarised in Table 1. Each parameter has an intuitive interpretation for its effect on generated patterns. The “fibre selection” field creates strands of collagen along the direction of fibres, a specific feature seen in some of the types of cardiac fibrosis we seek to recreate (Figure 1). Some parameters controlling this field are fixed at values judged appropriate for this specific application.

**Table 1:**
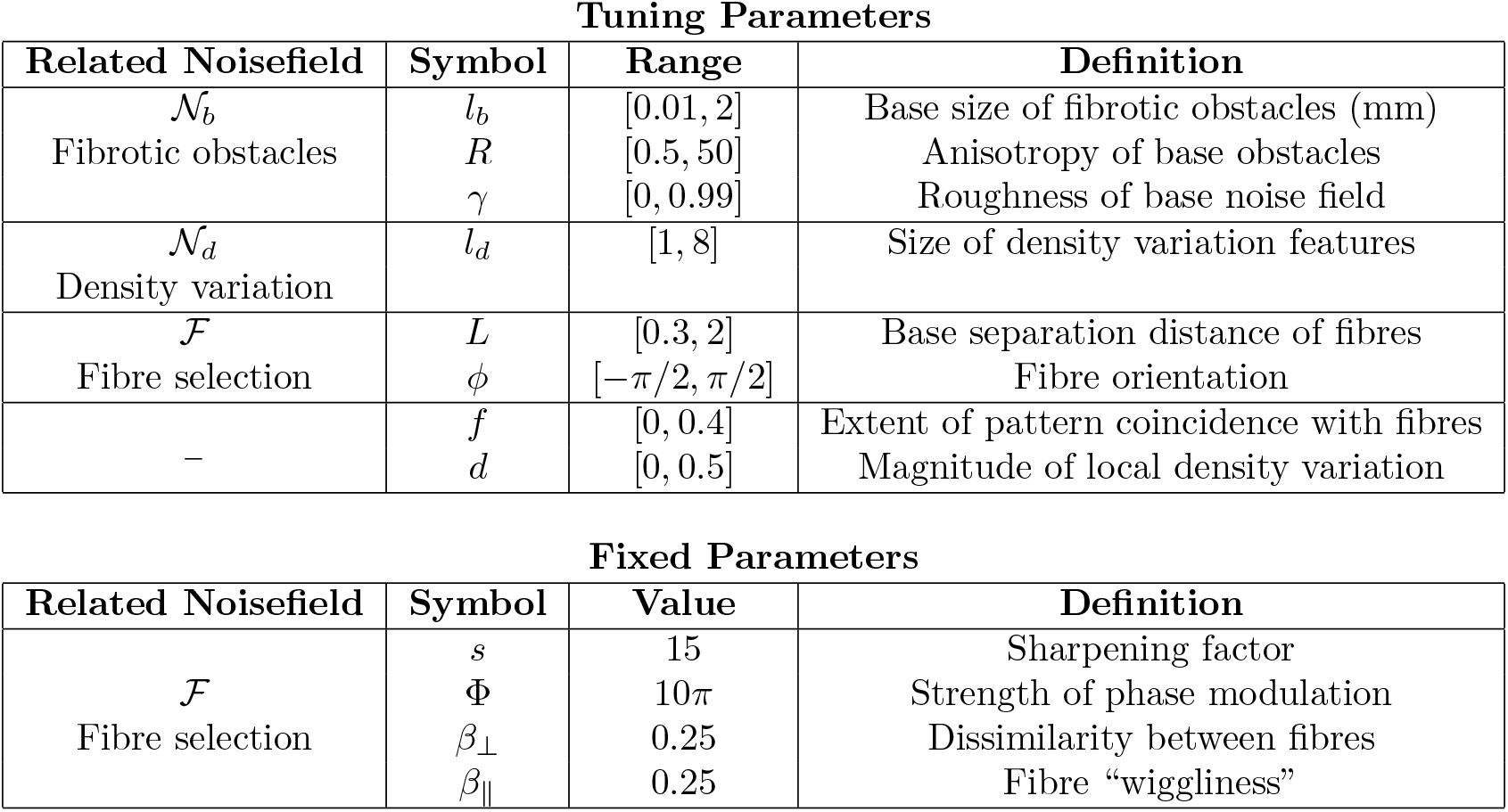
Tuning parameters of the generator function. Also listed are the noisefields that each parameter affects, and an intuitive understanding of their effects on generated patterns. Variable parameters are those that were varied to capture the different types of patterning considered in this work, and were searched over the quoted ratios during automated generator tuning. Note that as *R* is a ratio, ln(*R*) was instead the varied parameter so that it could still be varied uniformly (see Methods section related to SMC-ABC).

**Tuning Parameters**
| Related Noisefield | Symbol | Range | Definition |
| --- | --- | --- | --- |
| $\mathcal{N}_b$<br>Fibrotic obstacles | $l_b$ | [0.01, 2] | Base size of fibrotic obstacles (mm) |
| | $R$ | [0.5, 50] | Anisotropy of base obstacles |
| | $\gamma$ | [0, 0.99] | Roughness of base noise field |
| $\mathcal{N}_d$<br>Density variation | $l_d$ | [1, 8] | Size of density variation features |
| $\mathcal{F}$<br>Fibre selection | $L$ | [0.3, 2] | Base separation distance of fibres |
| | $\phi$ | $[-\pi/2, \pi/2]$ | Fibre orientation |
| – | $f$ | [0, 0.4] | Extent of pattern coincidence with fibres |
| | $d$ | [0, 0.5] | Magnitude of local density variation |

Table 1: Tuning parameters of the generator function.
| Related Noisefield | Symbol | Value | Definition |
| --- | --- | --- | --- |
| $\mathcal{F}$<br>Fibre selection | $s$ | 15 | Sharpening factor |
| | $\Phi$ | $10\pi$ | Strength of phase modulation |
| | $\beta_\perp$ | 0.25 | Dissimilarity between fibres |
| | $\beta_\parallel$ | 0.25 | Fibre “wiggleness” |

The remaining parameters of the generator are allowed to vary, allowing generation of a wide variety of patterns. We refer to these parameters as “tuning” parameters, and use our automated tuning process to select their values in response to the provided “target” patterns. Figure 2 shows a range of example patterns that can be produced by the generator, and the constituent noisefields used in their creation. Many different patterns can be produced by varying the tuning parameters, which include both the properties of the underlying fields and the parameters controlling their weighted combination. Our use of automatic threshold selection (see Methods) allows for the creation of patterns of any collagen density. The irregularity and wandering direction of the “fibres” in the fibre selecting field may appear striking, but it should be remembered that this field acts to place fibrosis between fibres, and indeed the patterns produced in our subsequent results are compelling. The extent of irregularity in the fibre selection field is easily varied by adjusting the parameters controlling its phase modulation, **Φ**, *β*_⊥_ and *β*_∥_ (Table 1).

**Figure 2:**
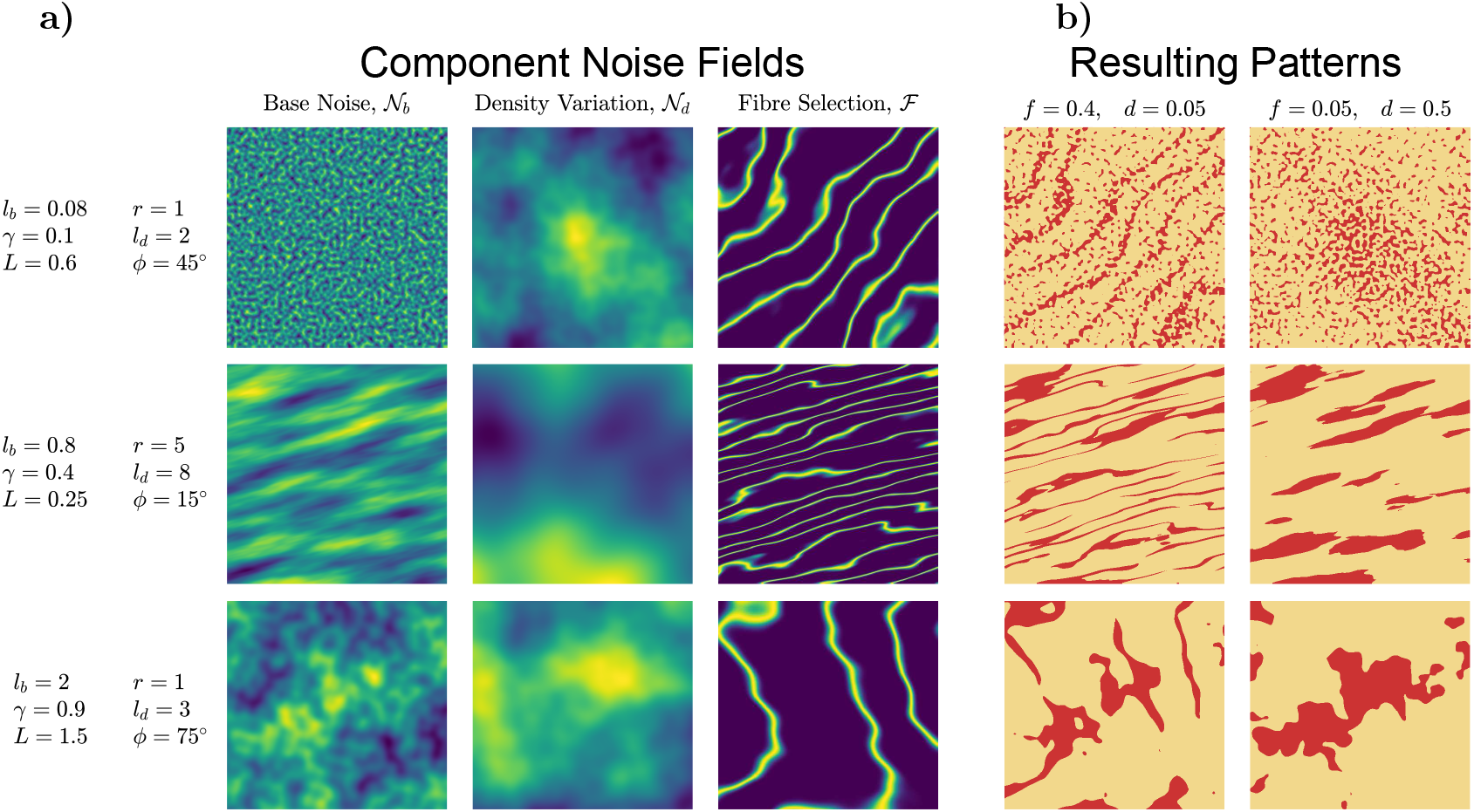
Example patterns created by the generator. **a)** The three noisefields that combine to create patterns of fibrosis. Each row shows realisations of the three noisefields for a different set of generator patterns, specifically a small feature size (top), high anisotropy and small fibre spacing (middle), and a large feature size with high roughness and large fibre spacing (bottom). Values for the generator parameters are given on the left of each row, with parameter definitions and the values of fixed parameters given in Table 1. **b)** The final binary patterns that result for different combinations of the three noisefields, as defined by fibreness *f* and density variation *d*, using a threshold chosen to obtain a collagen density of 20%. Pictured are a pattern with strong contribution of the fibre-selecting field and weak density variation (first column), and a pattern with weak contribution of the fibre-selecting field and strong density variation (second column). The values of *f* and *d* defining these choices are presented above the columns.

### Matching to Pattern Metrics Enables Automated Generator Tuning

#### Recovery of Target Patterns

We create a novel set of pattern quantification metrics, that then serve as low-dimensional representations of the most important pattern features, or in the ABC context, as our summary statistics [24]. These metrics use the power spectrum derived from the Fourier transform (full details in Methods). Before advancing to recovery of the histological patterns, we first demonstrate the efficacy of these metrics and sequential Monte Carlo approximate Bayesian computation (SMC-ABC) for automatic tuning by creating populations of patterns matched to three artificial target patterns, each produced using our generator with a unique set of parameter values. Targets created by our generator are guaranteed to fall within the space of patterns that it can recreate, allowing the automated tuning to be tested using targets that are known to be reachable. This approach also allows us to consider the recovery of the parameter values that were used in generating the target patterns.

The SMC-ABC algorithm with 2000 particles was applied to each of the three targets (Figure 3), generating sets of matching patterns. This relatively large number of particles was chosen to better ensure the method properly explored the parameter space, however runs with far fewer particles were also seen to successfully converge towards low-discrepancy populations. The method was run with a very strict target discrepancy, so that termination occurred when improvement of the population discrepancy became too slow (details in Methods). Upon termination, the set of output particles is a sample from the ABC-approximated posterior associated with the imposed statistical model (see Methods), but associated with each particle is also a pattern with low discrepancy from the target pattern. Therefore, after discarding the duplicated particles, the population of associated patterns can be used directly as a set of new realisations of the patterning in the target image. In order to demonstrate the suitability of these populations on the whole (as opposed to, for example, presenting only the lowest-discrepancy patterns or hand-choosing the best visual matches), all visualised samples from populations in this work use the patterns corresponding to the 0^th^, 20^th^, 40^th^, 60^th^, 80^th^ and 100^th^ quantile of discrepancy value.

**Figure 3:**
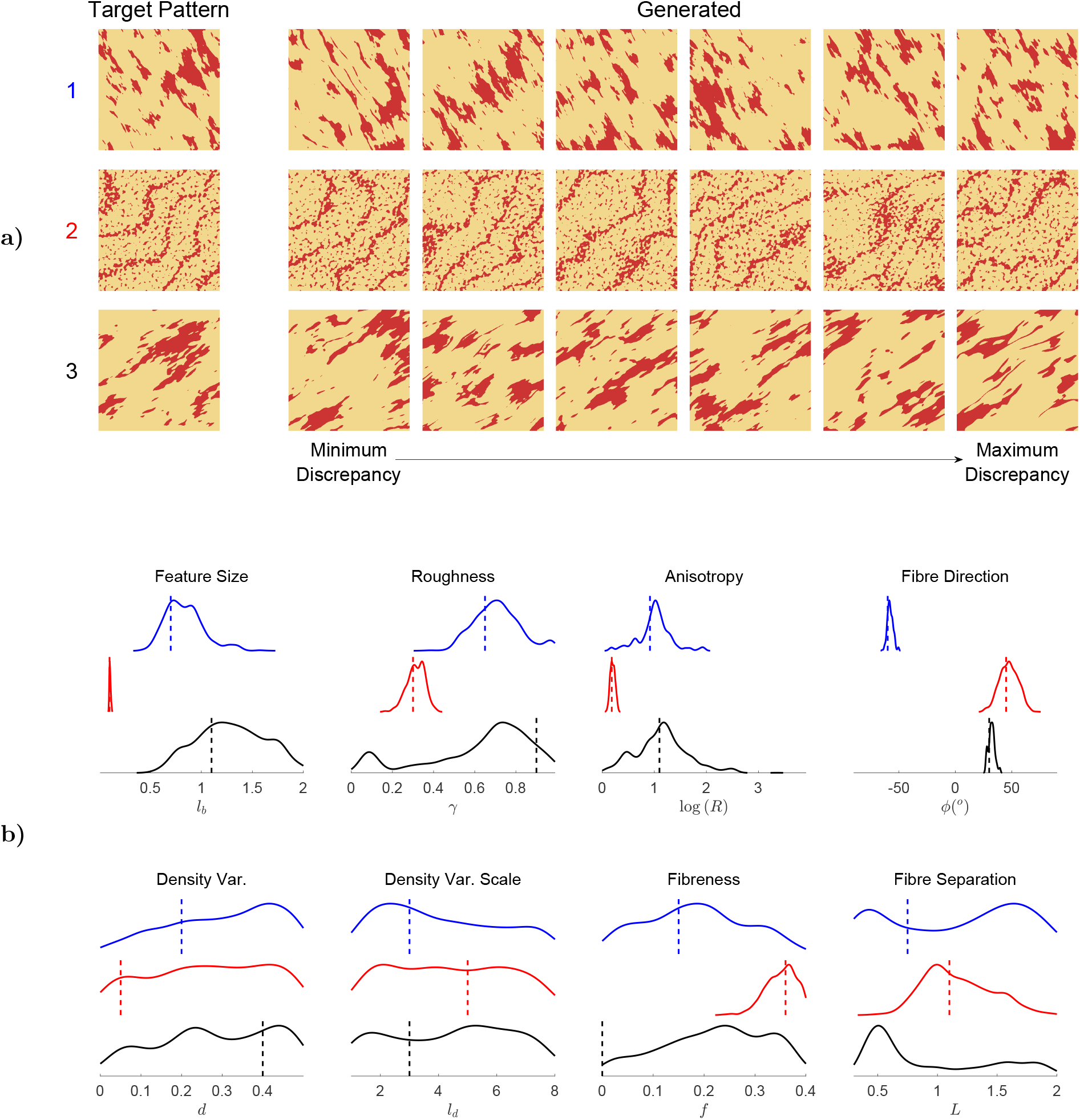
Successful pattern generation via power spectrum metrics and SMC-ABC optimisation. **a)** Samples from the sets of patterns obtained in matching three different target patterns, using our selected metrics and SMC-ABC. Visualised are the patterns selected by quantile (see text), with discrepancy increasing from left to right. For all target patterns, a great deal of visual similarity is achieved. **b)** Marginal distributions of the generator parameter values for the populations matched to the three target patterns (blue, red, and black solid lines, respectively), along with the original values used to generate the target pattern (vertical dotted lines). The parameters controlling the base noise field 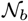 (top row) are all selected correctly and typically with high specificity, indicating a successful encoding of these properties by the pattern metrics. On the other hand, the parameters controlling the other two noisefields (bottom row) are typically estimated with low specificity, although a strong fibrous component to the target pattern does result in a successful recovery of these properties in the generated population. Given the good matching demonstrated in **a)**, parameters that are not estimated to high specificity can simply be interpreted as of less importance to achieving matching patterns in those cases.

Samples from the populations matched to each of the three target patterns are presented in Figure 3a. For all of the patterns, the sizes and orientations of collagenous obstacles are consistent between the target and generated patterns. Notably, even when both fibrous collagen strands and diffuse obstacles are present (pattern two), this type of patterning is recovered in all of the generated patterns. We stress, however, that the *positioning* of fibres and obstructions in the generated patterns differs from the target patterns. That is to say, as desired, our methodology is not simply producing near-copies of the target image, but instead producing new realisations of the same patterning that underlies it. Matching this patterning is achieved solely by minimising discrepancy in terms of our metrics, indicating their ability to encode important pattern properties.

In a small number of the generated patterns, minor differences with the target image can be seen. For the first and third target patterns, long, thin strands of collagen occasionally appear in generated patterns, but are not present in the target. The second target pattern shows a relatively even density of collagen deposition throughout the pattern, but a proportion of the generated patterns show (typically minor) spatial variance in collagen density. This occurs because our selected metrics describe patterns in terms of a set of basic features, and thus there is the potential for patterns that are similar in terms of these basic features to differ when considered in the full-dimensional space (that is, visually). For example, fibrous collagen strands can be an alternative means of achieving the target pattern’s anisotropy, and thus sometimes appear in the generated patterns even when not present in the target. The occasional pattern showing minor imperfections is not a significant issue, as once a population of matching patterns has been generated, it can then be used to select satisfactory tunings of the generator. We demonstrate this for the histological patterns of cardiac fibrosis we match to subsequently.

#### Recovery of Generator Parameters

Using artificial target patterns also allows us to consider how the parameter values of the pattern sets selected by SMC-ABC compare to the parameter values that were used in the generation of the target patterns. Under the Bayesian paradigm, a parameter “estimated” with high precision (low variance) implies that the value of that parameter is of high importance to the generation of low-discrepancy patterns. On the other hand, parameters estimated with low precision do not have a significant effect on the pattern metrics. The goal of SMC-ABC is therefore not to recover all of the parameter values used to generate the target patterns to high specificity, but instead to observe posterior distributions that are consistent with those parameter values, and then use the variance in values of the different parameters to understand their relative importance. It is only a concern if the metrics imply a parameter is unimportant, but it does actually have a significant effect on the resultant patterning.

Figure 3b shows the marginal distributions of the parameters selected across the populations of generated patterns (including duplicates in this case, for a correct posterior sample). In the populations matched to all three targets, the feature size *l*_*b*_, anisotropy ratio *R*, fibre orientation *ϕ* and to a slightly lesser extent the roughness, *γ*, are all estimated to match the parameters used to generate the target, and with relatively high precision. This indicates the successful encoding of all of the properties affecting the most important noisefield (the base noisefield 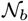) by the FFT-derived metrics. On the other hand, the density variation field’s spatial scale, *l*_*d*_, and strength, *d*, take on values across the whole prior, implying that the properties of this field are judged to be less important by the pattern metrics. For the most part this is correct, given the good visual fidelity of the generated patterns (Figure 3a) regardless of the value of *d* selected. However, it also allows large *d* values to be selected in matching to the second target pattern, where the target pattern has low *d* and the difference sometimes becomes apparent in the generated patterns. Importantly, when we consider the histological patterns of microfibrosis in the following section, our metrics do correctly bias towards selecting high *d* values in matching the targets that exhibit it.

The selection of parameter values controlling the fibre-selecting field depended strongly on the target pattern. When the fibre-derived component had a moderate or high contribution to the target, this contribution was correctly estimated. In the case of pattern two, where the contribution of this field is strong and fibres are obvious in the target image, the specificity of this estimation was also high, and the fibre spacing was also well estimated. However, for the third target pattern with no contribution of the fibre-selecting field (*f* = 0), many of the generated patterns still feature *f* values towards the upper end of the range of its values. This is likely the cause of the second mode in the distribution of roughness values for this population, as additional complexity that would be added to the generated patterns by a large roughness value is instead being created by fine fibres present in the patterns with high *f* values.

On the whole, our metrics derived from the power spectrum effectively encode the key properties of patterns, with very good visual agreement between the target patterns and generated patterns with low discrepancy. SMC-ABC successfully generates populations of low-discrepancy, automatically finding tunings of the generator that produce new realisations of a provided target patterning. Generator tunings used to create the target data were for the most part also recovered by the SMC-ABC process, with the parameters that were estimated with low precision also tending to show little impact on the generated patterns. With our automated tuning via pattern metrics validated, we now go on to demonstrate that our generator can produce the types of patterning seen in histologically-observed cardiac fibrosis.

### Perlin Noise Successfully Generates Patterns Representative of Histologically Observed Fibrosis

We apply our automated tuning procedure to create patterns representative of the different types of microstructure in cardiac fibrosis (Figure 1). Again using 2000 particles, SMC-ABC was run until particle degeneracy implied a lack of population improvement, resulting in populations of between 900 and 1000 unique, representative patterns for each class of fibrosis. These populations possess low discrepancies from the target pattern in terms of our metrics (see Figure S1), and capture the key properties of each class of fibrosis (Figure 4). For example, the generated patterns of interstitial fibrosis correctly consist of long strands of collagen deposited along obvious, but slightly disorganised fibres. In addition to a successful matching to the patterns’ distinguishing features, the generated patterns also display the correct amount of density variation (evident in the diffuse and patchy patterns, and absent in the interstitial and compact patterns).

**Figure 4:**
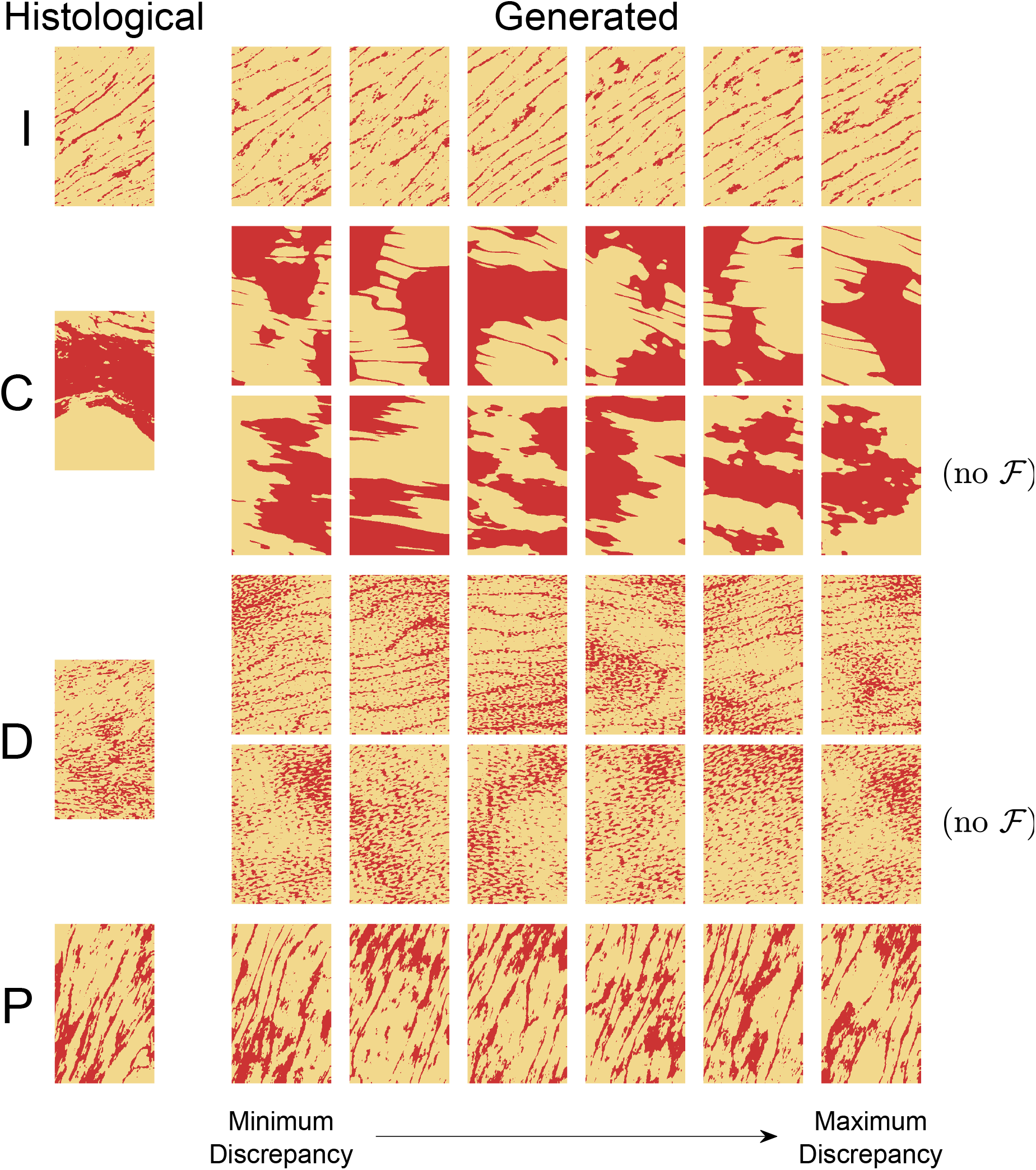
Successful generation of the different types of microfibrotic patterning. The target histological images (leftmost column) for interstitial (I), compact (C), diffuse (D) and patchy (P) fibrosis, and a sample of the patterns output by our generator after SMC-ABC tuning process (remaining columns). The generated patterns capture the distinguishing features of the four types of microfibrosis — long fibre-aligned stands for interstitial, large contiguous clumps for compact, speckles of varying density for diffuse and anisotropic clumps with fibrous influence for patchy. Generated patterns for compact and diffuse fibrosis include both those attained via the standard SMC-ABC tuning, and a tuning with the fibre-selecting field turned off (the rows marked “no 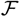”).

The generated patterns for compact fibrosis display large regions wholly occupied by collagen, and for diffuse fibrosis tortuous paths through regions occupied by many small obstacles, as desired. In both cases, however, the SMC-ABC tuning using our metrics consistently results in the inclusion of strands of collagen along fibres that are not present in the target histological images for these patternings. Including a fibre-selecting effect (*f* > 0) allows the generated patterns to better resolve subtle anisotropies in the target images (made evident by the fibre direction consistently being selected to match the direction of greatest anisotropy in the target image), but at the cost of pattern fidelity. For this reason, we also created populations of patterns of diffuse and compact fibrosis using SMC-ABC but with a reduced set of parameters, fixing the contribution of the fibre-selecting field at *f* = 0 (and *L* now rendered irrelevant). These generated populations, also presented in Figure 4, still capture all of the key pattern features but now without the presence of fibrous collagen strands. In fact, the shapes of the clumps in compact fibrosis now also show rougher edges, in better agreement with the histological example.

We have used SMC-ABC as an automated tuning process for finding optimal patterns in terms of metric discrepancy. However, the selections of parameter values across the matched populations (Figure S2) are also in accordance with their practical interpretations presented in Table 1, demonstrating the potential for manual tuning by the user according to these properties. Feature sizes were selected to match those evident in the histological images (very small for diffuse, moderate for interstitial and patchy, large for compact), and roughness values according to the complexity of feature shapes (lowest for the rounded shapes in diffuse fibrosis, highest for the rough edges in interstitial and patchy fibrosis). Selected fibre orientations consistently matched the anisotropies in the target images, and the populations for the fibrosis types with the most obvious density variation (diffuse and patchy) showed the strongest bias towards high contributions of the density variation noisefield.

We also now use our matched populations to derive single sets of parameter values that we consider representative of the different types of fibrosis. This both allows the generation of further realisations without a user needing to run the tuning algorithm, and demonstrates the ability to select only a single set of validated parameter values and still create multiple realisations of a desired patterning. Representative sets of parameters were calculated from the SMC-ABC populations by taking the modes of each parameter’s marginal distribution in the population (as estimated by MATLAB’s kernel density estimation function ksdensity). This resulted in the parameter values given in Table 2. The parameter sets in Table 2 generate low-discrepancy patterns that match very well visually with the histological images (Figure S3), each still with its own unique placement of obstacles.

**Table 2:** Generator tunings for creation of new microfibrosis realisations. The modes of the marginal distributions of the generated populations of representative patterns, suggested as values for the generator’s parameters that create new realisations of the desired type of microfibrotic patterning. Fibre orientations, *ϕ*, and density were selected to match the histological images considered in this work, but can be varied as desired by the user.

| Pattern | $l_b$ | $\gamma$ | $R$ | $\phi$ | $d$ | $l_d$ | $f$ | $L$ | Density |
| --- | --- | --- | --- | --- | --- | --- | --- | --- | --- |
| Interstitial | 0.24 | 0.96 | 1.89 | $34^\circ$ | 0.32 | 4.67 | 0.30 | 0.31 | 0.096 |
| Compact | 0.96 | 0.59 | 2.47 | $-9^\circ$ | 0.44 | 2.03 | 0 | - | 0.472 |
| Diffuse | 0.07 | 0.44 | 2.17 | $11^\circ$ | 0.49 | 1.22 | 0 | - | 0.220 |
| Patchy | 0.32 | 0.78 | 2.50 | $68^\circ$ | 0.42 | 2.10 | 0.38 | 0.31 | 0.269 |

### Minor Variations in Microfibrotic Structure can have Profound Electrophysiological Impacts

In the previous sections, we have presented and validated a powerful tool for the automated, computational generation of new realisations of fibrotic patterning. We now demonstrate the importance of this tool, by presenting simulations that highlight the effects of microscopic variability on the electrophysiological impact of cardiac fibrosis. Our generator allows this variability to be investigated without the need for large amounts of invasive, *ex vivo* collection of microfibrosis imaging data, by rapidly generating new realisations with little compromise of physiological accuracy and a greater degree of control.

Electrophysiological impact is explored by stimulating one side of two-dimensional slices of myocardial tissue, into which the patterns of collagenous obstruction are placed (Figure S5). Simulations are performed under conditions of chronic atrial fibrillation (cAF), including fast pacing, given the significant evidence linking fibrosis to the sustenance and post-treatment reoccurence of this condition [25, 26]. Impact is judged in terms of two markers of arrhythmogenic risk, activation delay (AD) [10] and unidirectional block [27]. If one pattern of fibrotic obstruction shows the potential for unidirectional block and another does not, this represents a significant difference in the potential to serve as substrates for micro re-entry, an arrhythmia precursor [7].

Figure 5 shows maps of AD for some of the patterns presented in Figure 4, using those generated without a fibrous component (no 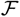) for compact and diffuse fibrosis. The corresponding activation time maps are provided in Figure S4. The left sides of the images, where the tissue is stimulated, show slightly negative AD values. This is because the fibrotic region causes a reduction in electrotonic (diffusive) loss of potential, and thus the wave of excitation propagates slightly faster. The other side of the fibrotic region consistently shows positive AD values, representing the delay in excitation propagation caused by navigating through and/or around the collagenous obstructions. The different classes of fibrosis show significantly different amounts of AD, but we note that some of this is attributable to the interplay between the different fibre directions associated with each and the fixed placement of the fibrotic region relative to the stimulus location. On the other hand, differences in AD within a single class of fibrosis owe solely to the effects of the precise arrangements of collagen in each pattern.

**Figure 5:**
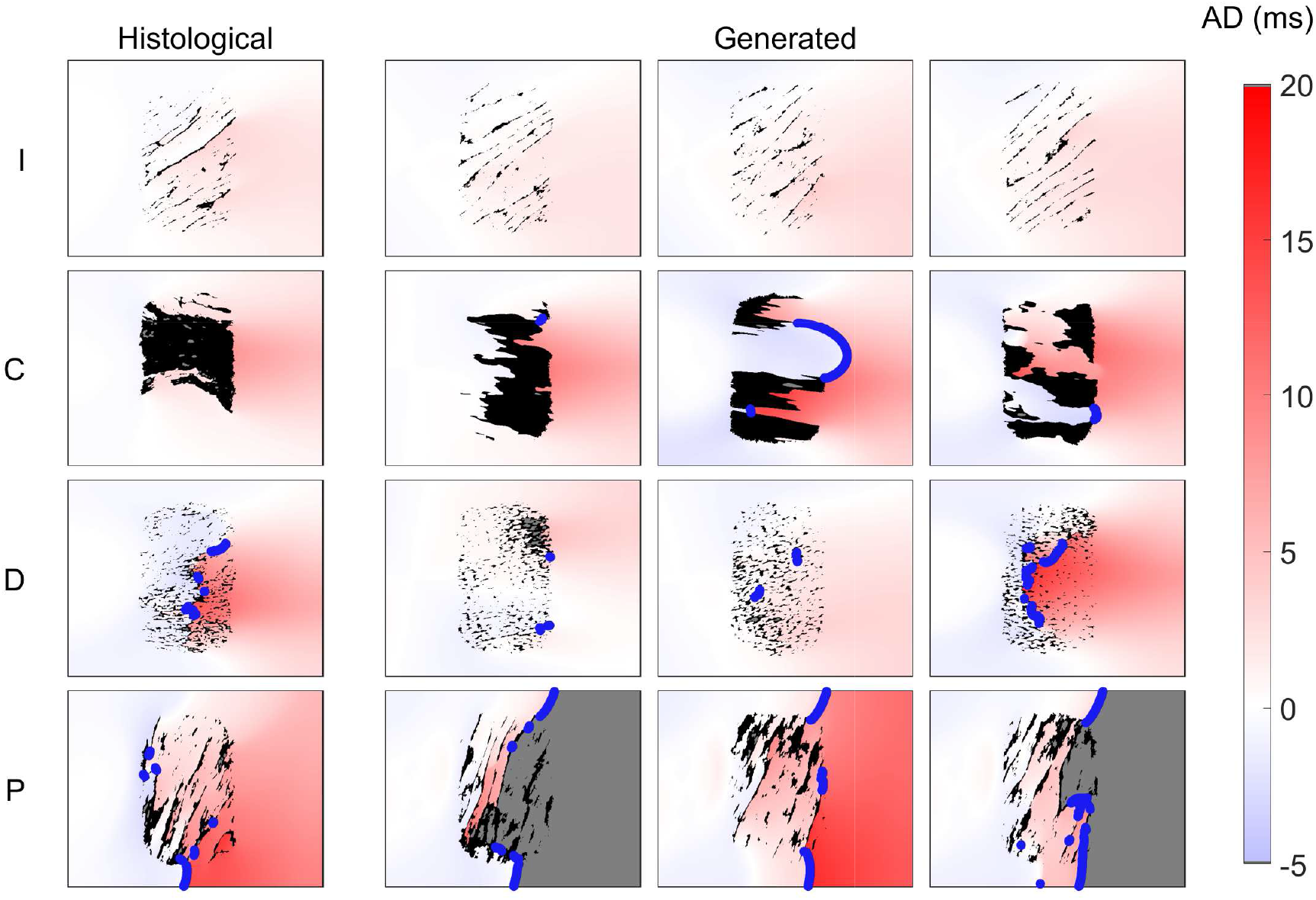
Activation delay (AD) arising due to different organisations of fibrosis under fast pacing and chronic atrial fibrillation conditions. Shown are maps of AD on 4.78 mm×3.68 mm slices of myocardial tissue, arising due to a fibrotic region of approximately 4.6 mm2 occupied with collagenous blockage corresponding to interstitial (I), compact (C), diffuse (D) or patchy (P) fibrosis. AD maps for arrangements of collagen derived from histology (left column) and produced by our generator (remaining columns) are shown. Delay is measured by subtracting activation time for the case with fibrosis present from activation time for the case with equivalent pacing conditions and fibre orientation, but no fibrosis present. Grey sites are those not activated by the stimulus pulse at all. In each case, the stimulus from the pacing regime that produced the most electrophysiologically notable activation pattern is visualised (from highest to lowest priority: complete conduction block, localised conduction block, maximum AD). Strong blue lines mark events of conduction block causing significant delay. The severity of activation delay is seen to depend strongly on the precise positioning of collagenous obstacles, even for the same type of fibrosis.

The pattern of AD due to interstitial fibrosis is wholly consistent between the histological pattern, and each of the generated patterns. This demonstrates the efficacy of the generator in creating physiologically realistic patterns that also behave equivalently, but does also suggest that the precise arrangement of individual obstacles is less important in this type of fibrosis. For the other three classes of microscopic patterning, however, significant differences are seen between the different pattern realisations. For compact fibrosis, when pathways through the fibrotic region do exist they have the strong tendency to block, with signal failing to propagate beyond the end of the pathway (due to the source/sink mismatch effect [2]). Unidirectional block in thin channels through infarcted regions has been established as a trigger for arrhythmia both experimentally [28] and computationally [29].

Diffuse and patchy fibrosis show far less predictable effects on excitation propagation. The histological example and one generated realisation (rightmost in Figure 5) of diffuse fibrosis exhibit enough block of conduction to force signal to wrap around and invade the fibrotic region from the far side from the stimulus location, while the other realisations show only minor disruptions to conduction and AD. There do not appear to be any obvious features of the patterning that could predict this significant deviation in activation behaviour, highlighting the importance of further study.

Patchy fibrosis shows a strong tendency to block above and below the fibrotic region, in part due to the weak propagation of the wavefront that moves in a direction mostly transverse to the fibres. However, whether or not propagation through the fibrotic region succeeds depends on its microscopic makeup. In the cases where excitation fails to propagate through the fibrotic region, there is a complete failure to activate the area beyond it. Examination of successive stimuli reveals further complexities to the activation behaviour of these two patterns, with one showing 2:1 block while the other cycles between complete block, partial block and normal propagation (see supplemental movies). Again, this clearly demonstrates the need for a more sophisticated understanding of how micro-scale arrangement of non-conducting protein deposits modulates the risk of arrhythmia associated with fibrosis-afflicted tissue.

## Conclusions

We have presented a new methodology that uses Perlin noise to generate new realisations of fibrotic patterns, that capture the complex arrangements of collagenous obstruction observed histologically. Using our method of characterising key pattern features to enable automated tuning of our generator, we successfully produced new realisations of the four types of microfibrotic structure in cardiac fibrosis [5]. We demonstrated the ability to use singular sets of generator parameters to produce new realisations, presenting suggested tunings for these four types of cardiac fibrosis that can be used in future *in silico* studies. As our simulations have indicated, a fuller understanding of how variability in microscopic collagen distribution modulates electrophysical impact is critical, and the key novel impact of our methodology is to remove limitations posed by the availability of *ex vivo* data whilst still being sophisticated enough to produce physiologically realistic patterns that match what data is available.

Our use of a very flexible generator allows it to produce other patterns beyond the examples we have used to demonstrate it, and in particular suggests that it should be able to capture patterns that are intermediary between the different established classes of cardiac fibrosis considered here. Our metrics allow for the automated SMC-ABC approach to tuning the generator in response to a new type of patterning to be generated, but we re-iterate that the parameters themselves all have intuitive interpretations that facilitate adjustment by hand.

A natural extension of our technique would be the construction of physiologically accurate fibrotic maps on accurate organ anatomies in three-dimensions. Perlin noise extends without modification to three dimensions, and can be evaluated at any point in space, permitting irregular meshes. Macroscopic data of fibrotic burden (that can be collected non-invasively [30, 31]) could be used to inform the collagen density used in different regions, or the macroscopic arrangement of fibrosis could also be generated as realisations of Perlin noise. A question is instead how to generate patterns that remain continuous when the fibre direction varies across the pattern, as it will over the larger physical distances involved. One possibility is a curvilinear coordinate system that tracks the varying fibre direction across the heart [32]. Our pattern quantification metrics also generalise to three dimensions, but would also need to be interpreted in such a coordinate system, presumably requiring the non-uniform variation of FFT in order to calculate the power spectra [33]. If the generated fibrotic maps were to be used for simulation, an additional concern is computational cost, given the number of mesh elements required to resolve such microscopic detail on the organ scale. A method for tailoring finite element meshes to represent the impacts of fibrotic obstruction, without invoking the resolution required to explicitly resolve it, has been presented in two dimensions [34].

The potential of a fast, computational generator of physiologically realistic patterns is not limited to computational studies of variability. Perlin noise has been used as a rapid generator of “breast phantoms” [22], artificial realisations of breast imaging data used to perform virtual clinical trials or obtain sufficient data for machine learning without concern of cost or radiation dosage to patients [35]. Our methodology achieves the same goal in the context of (micro)fibrosis, and extends that work in that the tuning process directly uses available imaging data, capturing the evident patterning in a quantitative manner. This allowed us to explicitly reproduce significantly more complex patternings than those required for breast phantoms. Our novel metrics and the SMC-ABC tuning approach are also wholly distinct from the fibrosis context, suggesting their applicability to other types of physiological patterning. With the generator modified to suit the new context (if necessary), our presented methodology allows for the rapid generation of very large numbers of pattern realisations, offering massive sample size extension to bolster machine learning methods that have continued to proliferate in the medical context [36, 37].

One minor limitation of our study is that we tuned fibre-selecting field specifically for the generation of collagenous strands aligned with fibres, and did not explore the effects of varying these parameters, nor the ability for SMC-ABC to automatically tune them. This might be relevant if other experimental data, inside or outside the cardiac fibrosis context, showed a fibrous component of a different character to the histological images we have considered here. Our chosen metrics were not always able to detect the presence or absence of fibrous strands, however this was easily resolved by first pre-selecting whether to include the field that generates these before performing the automated tuning process.

In the heart, arguably the best-understood organ from both a physiological and mathematical perspective, the importance of fibrosis microstructure has already been identified as a key issue [38, 7]. Variability on the microscopic scale has been explored computationally in terms of density of obstruction for diffuse fibrosis [39, 40], including the differential effects on excitation propagating in directions longitudinal or transverse to the fibre direction [41]. A recent work incorporated interstitial and diffuse fibrosis into a larger-scale model, using image data to inform density but without an attempt to explicitly capture the precise microfibrotic structure [17]. Our work, however, is the first effort to explicitly consider micro-scale variability using histologically-verified patterns, and demonstrates a powerful framework for further such research towards understanding the impacts of fibrosis. This is extensible to pulmonary and hepatic fibrosis where extracellular matrix structure is strongly implicated in the progression of these diseases on a cellular level [42, 43].

## Methods

### Histological Patternings of Microfibrosis Structure

We use as reference images the histological sections of human heart epithelial layers collected by Kawara *et al.* [10], and presented by de Jong *et al.* [5] as examples of the four primary types of cardiac fibrosis. We simplify these images into patterns of the presence or absence of collagenous obstruction (Figure 1), in order to clearly demarcate conductive and non-conductive regions for subsequent simulation. However, we note that our metrics for pattern quantification do not depend upon the binary nature of the data and could be expected to also successfully quantify “continuous”, non-thresholded image data.

Thresholding of the histological images used the green channel (*G* < 180 indicating collagen), as a simple method for distinguishing the red regions marking collagen deposits from the yellow regions marking myocardial tissue in the original images. Representative 250 × 400 pixel sections of each image were selected to ensure a consistent data size between the different classes of fibrosis, and to speed up simulations. For compact fibrosis, the regions at the bottom of the image were also removed due to falling on the sample boundary.

### Metrics for Pattern Quantification

We consider patterns in terms of their fast Fourier transform (FFT), a common tool in image processing that expresses all of the information contained in an image in the frequency domain [44]. The frequency components of the FFT correspond to variation in the image at different spatial scales and along different directions, with the intensity (squared magnitude) of these components quantifying their relative importance in re-constructing the image. Thus the *power spectrum*, composed of the intensities of each of the Fourier components, can be interpreted as quantifying how fibrotic obstacles of different size and orientation contribute to the image. We use only the power spectra (and no phase information from the FFT) to quantify patterns, as we are not concerned with the precise positioning of fibrotic obstacles in the image, only their sizes, shapes and positions relative to one another.

The high dimensionality of the power spectra (*d* = *N*_pixels_) makes direct comparison of patterns difficult. Instead, we create much-lower dimensional vectors of pattern metrics that encode some of the most important information from the power spectra. Matching is then considered in this lower-dimensional space, in the same way that summary statistics are used in the ABC literature [24].

Power distributed across high frequency components indicates fine pattern structure, while a concentrated low frequency spectrum corresponds to large feature sizes. Moreover, the directionality of the spread of power corresponds to anisotropy in the pattern. We access this information from the noisy power spectra by first shifting them so that their lowest-frequency components are centred, then applying smoothing via a Gaussian blur (with standard deviation *σ* = 4 pixels). The contour lines (of equal power) in these smoothed power spectra are then approximately ellipsoidal, with axis lengths corresponding to the range of frequencies contributing to some power quantile. The orientation and eccentricity of these ellipses correspond to the orientation and extent of anisotropy in the pattern, respectively. Fine-scale effects will involve higher frequencies and thus only affect the contours of ellipses containing large proportions of the power in the spectrum. Thus, multiple ellipses corresponding to different power quantiles encode the pattern behaviour over multiple spatial scales, and so we use these ellipses for pattern quantification.

Specifically, we use the orientation, major axis length and minor axis length of the ellipses containing multiples of 10% from 10% to 90% of the total power in a power spectrum as our metrics. This results in 27-dimensional metric vectors consisting of this set of three properties for nine ellipses. Ellipses were generated by finding cutoff power values for the different proportions of total power (power quantiles), creating binary masks specifying the frequency components falling above this cutoff value. Best-fit ellipses to these masks were then generated using the regionprops function of MATLAB’s image processing toolbox.

Figure 6 presents the ellipses associated with the histological images of collagen patterning in Figure 1, demonstrating how they summarise the important properties of the patternings. The large contiguous regions in compact fibrosis are represented by very small ellipses (low frequencies sufficient to explain much of the pattern). Ellipse eccentricity encodes the anisotropy of the pattern, with high eccentricity seen in particular for interstitial fibrosis. The use of ellipses at different power levels successfully summarises the behaviour of the pattern over different scales. For example, the difference in eccentricity of the lowest-power ellipse between the interstitial and diffuse patterns indicates that at large scales, the interstitial pattern is anisotropic, whereas the diffuse pattern is essentially isotropic. Similarly, only the highest-power ellipse is eccentric for compact fibrosis, because anisotropy only becomes evident when considering the subtle details of what is essentially a contiguous and isotropic (rectangular) obstacle.

**Figure 6:**
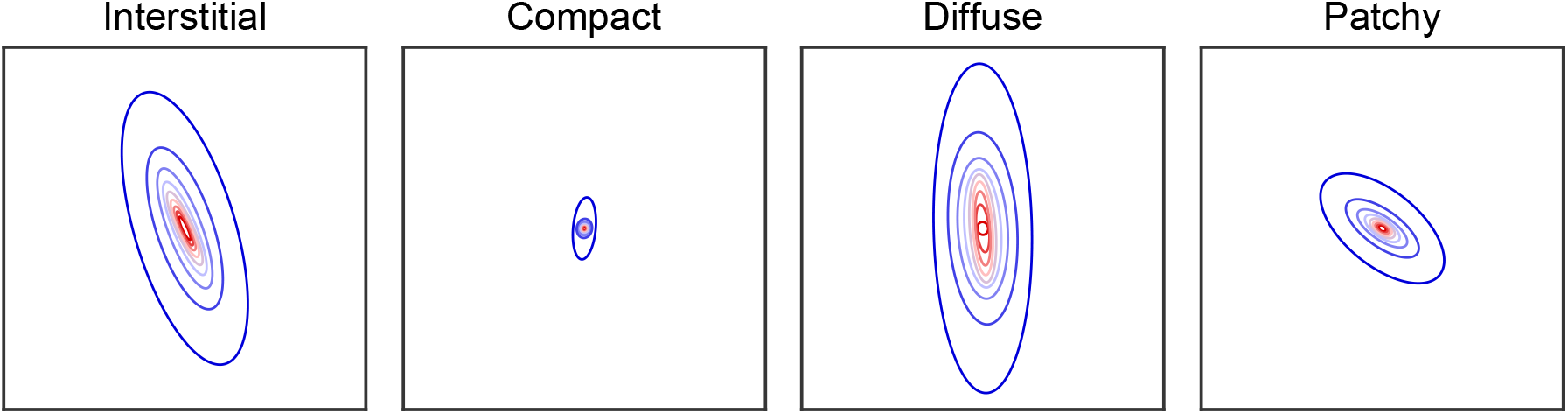
FFT-derived metrics for pattern quantification. Ellipses from the smoothed power spectra, corresponding to 10% (dark red) through to 90% (dark blue) of the total power, that serve to quantify the most important features of the histological images of fibrosis shown in Figure 1. The axes are consistent across the four patterns to allow for visual comparison, and unmarked because only scale is important. As described and demonstrated in the text, the different properties of each of the ellipses (length of major axis, eccentricity, orientation) successfully encode the key features of the different types of microfibrotic structure.

### Perlin Noise

Perlin noise is a computer graphics technique for the procedural generation of textures [21]. One of its primary appeals is that it generates patterns composed of ‘features’ (regions of similar value) of controllable size, thus serving as a powerful building block for the construction of a wide variety of patterns and textures (Figure 7). More formally, its power spectrum approximates a constant value across a band of frequencies (and zero outside of that band) [45]. This is in contrast to Gaussian random fields, another popular noise-generation technique that has non-zero power at all frequencies and thus produces less distinct features (Figure 7, vi). We present the details of the technique and our implementation in the supplementary material.

**Figure 7:**
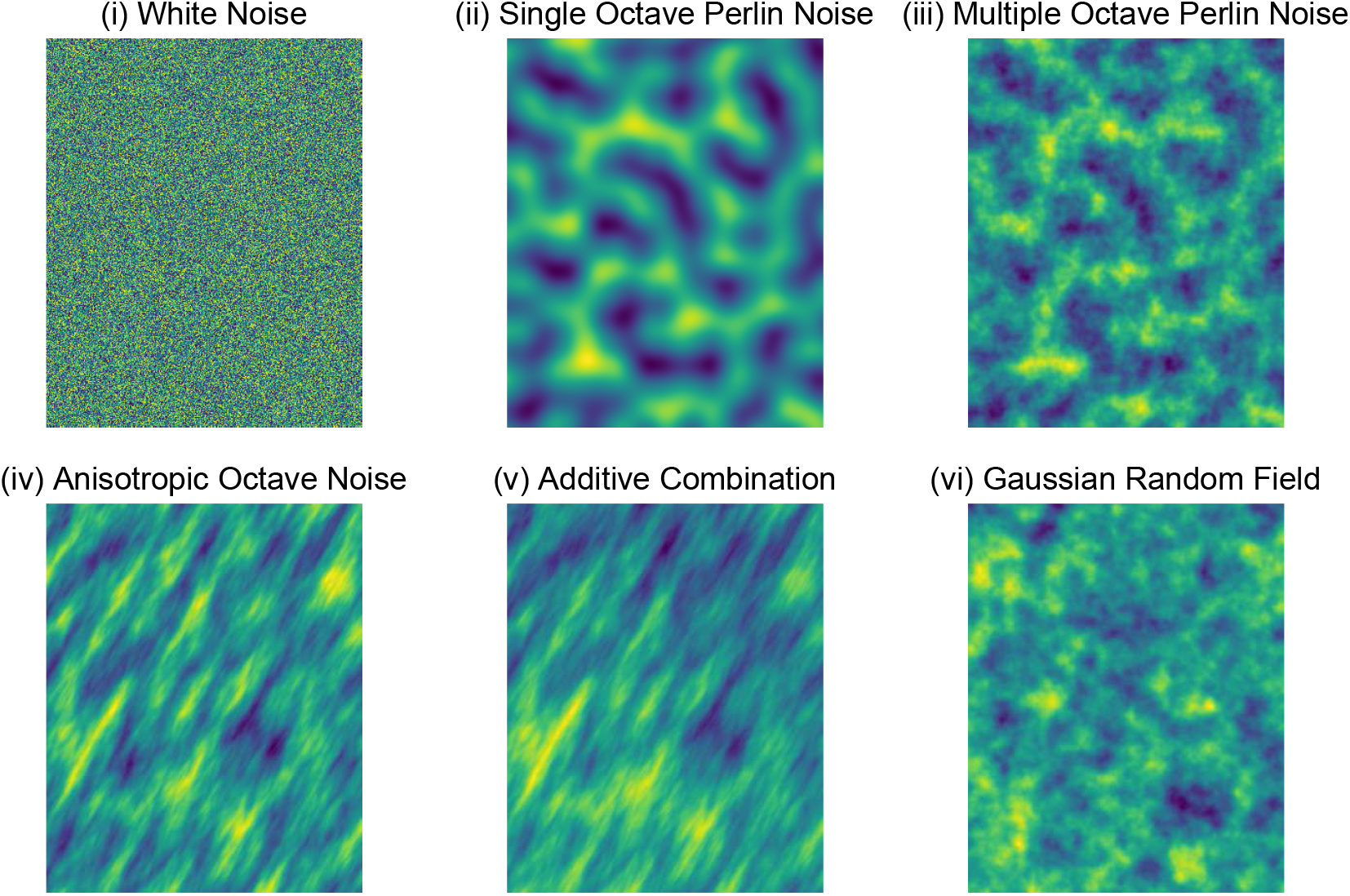
Example realisations of different noisefields. **(i)** White noise (uniform random value at each pixel). **(ii)** Single-octave Perlin noise. **(iii)** Refinement of the Perlin noise field in (ii) by additional octaves, as described in equation (1). **(iv)** Features with a specified anisotropy achieved via squeeze mapping and rotation of the input points, as described in equation (2). **(v)** Mixture of (iv) with an isotropic, large-feature Perlin noise field creates localised regions of generally higher or lower value (compare top and bottom of image). **(vi)** A Gaussian random field (with Matern-3/2 covariance function), for the sake of comparison. Full information regarding the generation of each of these patterns is provided in the supplementary material.

Traditional Perlin noise uses a hashing algorithm to select from a reduced pool of vectors, in order to avoid generating the large numbers of random vectors that it would otherwise require [21] (further details in supplement). The permutation tables used as part of this hashing operation can be generated from a single random seed, and new realisations of the noise with the same properties can be generated simply by changing this seed. This approach also aids reproducibility.

Following Ken Perlin’s original demonstration of his noise-generating technique [21], we combine multiple Perlin noise fields of doubling frequency, termed *octaves*. Successively downweighted, the higher-frequency fields become a series of refinements to the base Perlin noise field, adding complexity to the shapes of features (compare Figure 7, ii and iii). Denoting by *P* (**x**) the process of finding the value of a Perlin noise field at the point **x** (as described in the supplement), we combine the values of multiple such realisations *P*_*i*_ to create refined noise fields 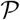 that are the Perlin noise we refer to in this work. These take the form

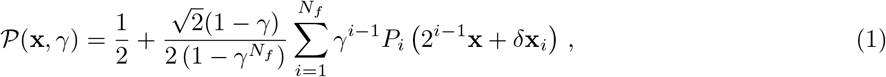

with *N*_*f*_ is the number of frequencies (octaves) that are combined (*N*_*f*_ = 4 in this work), and *γ* the intensity ratio successively applied as the frequency increases. We refer to *γ* as the “roughness” of the noisefield, as it controls the roughness of the shapes of its features. The *δ***x**_*i*_ are offset vectors with each element selected from the uniform distribution 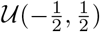, included to de-emphasise grid effects. Our definition of Perlin noise uses a positive shift of a half as well as a scaling factor to make the range of the noisefield [0, 1] (the original range of single-octave Perlin noise in two dimensions is 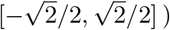.

Equation (1) generates continuous patterns with isotropic features of unit size, due to the unit grid spacing used by the base Perlin noise. In order to obtain features of a desired size and anisotropy, the set of evaluation points must first be transformed relative to the Perlin noise grids. For example, in order to double feature size, the set of input points should be scaled by a half, so that they will lie closer to one another relative to the fixed sets of gridpoints. Features of a desired size, orientation and anisotropy are obtained by application of a scaling, a rotation, and then a squeeze mapping. That is, in two dimensions a noise field 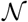 with features of overall size *l*, anisotropy ratio *R* oriented in direction *ϕ*, and roughness *γ* may be generated using

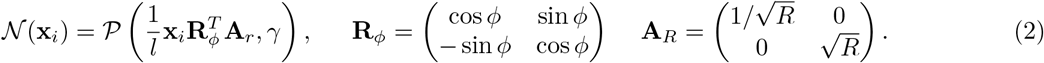

The notation **x**_*i*_ = (*x*_*i*_, *y*_*i*_) is given to evaluation points, in anticipation of evaluation of the noisefield at a large number of points in order to create patterns.

Denoting **X** the *N* × 2 matrix of points at which the noisefield is to be evaluated, each **x**_*i*_ is then a single row of this matrix. In this case, **X** is the set of co-ordinates of the pixels in the pattern images that we create. However, the same approach applies even when the set of evaluation points is irregular (consider, for example, physiologically accurate heart anatomies), and could be used in three dimensions via standard extensions of the rotation and squeeze mapping operations.

### Fibrotic Pattern Generation via Perlin Noise

We use a combination of Perlin noises of form (1) to generate images with continuous-valued pixels, which are then thresholded to become binary patterns representing the presence or absence of fibrotic obstruction. We denote a generated image 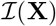, with **X** the set of pixel coordinates. Pixel coordinates are expressed in terms of physical units (mm), with the pixel width of 1/136 mm estimated using a scale bar from the histological images (see [5]). The generator is best described in terms of a base generator, 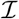, that produces patterns with a fibre orientation of 0° (from the horizontal). A rotation of its input points then generates patterns of the desired orientation *ϕ*, 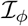. The generator takes the form

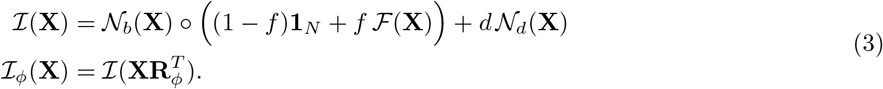

Here the notation 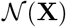 refers to the (column) vector generated by evaluating the noisefield 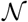 at each of the points in **X**, **1**_*N*_ = (1, 1, …, 1)^*T*^ is a vector of *N* ones and *◦* denotes the Hadamard (element-wise) product 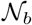 and 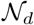 are two separate Perlin noise fields, and 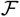 is a “fibre-selecting” field that also uses a Perlin noise field in its definition. The forms of these fields are given subsequently. The parameters *f* and *d* control how these three constituent fields are combined to produce the final image. Specifically, *f* controls the extent of collagen deposition along fibres (as seen for example in interstitial fibrosis), and *d* controls the extent the fibrotic density varies across different regions of the pattern. The remainder of the generator’s parameters lie within the definitions of the component noise fields that we now describe, and also listed in Table 1.

The base Perlin noise can be expressed in element-wise form as

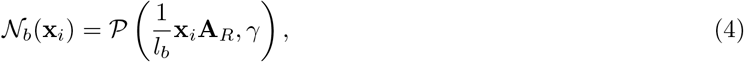

and does the primary work in creating the fibrotic obstacles. The transformation (2) is used to specify the size and shape (anisotropy) of these obstacles. These properties vary between the different patterns, and so the feature size *l*_*b*_, roughness *γ* and extent of anisotropy *R* are left as tuning parameters of the generator.

The density variation Perlin noise (also in element-wise form) takes the form

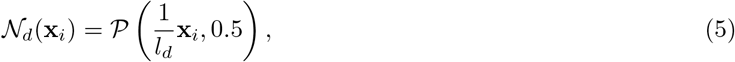

using large, isotropic features with somewhat low roughness (0.5) to create broad and relatively smooth regions. When this noise is additively mixed into the generator, it creates localised regions biased towards higher or lower intensity values (Figure 7, (v)). After thresholding, these shifts act to adjust the density of fibrosis in those regions, capturing the variation in local density seen in particular in the images for diffuse and patchy fibrosis (Figure 1). The spatial scale of these regions of density variation is a tuning parameter, *l*_*d*_, and the overall extent of density variation is controlled by *d*, as described above.

The fibre-selecting pattern

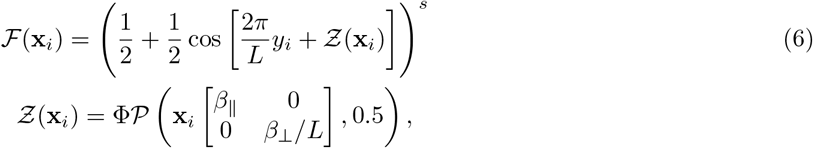

uses a sinusoid transformed to take values in the range [0, 1]. Horizontal fibres are generated by evaluating at only the *y* components of the input points, with the frequency 2π/*L* creating a fibre spacing of *L*. A large exponent (here termed a sharpening factor, *s*) is used to only retain values close to one, converting the sinusoidal pattern to a set of regularly spaced ridges that serve as thin fibres.

The phase of the sinusoid is also varied randomly through the image by a phase modulation field, 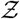, introducing natural irregularities into the shape, thickness and spacing of fibres. This field is also generated using Perlin noise, with Φ controlling the overall strength of the phase modulation effect, and *β*_⊥_ and *β*_∥_ controlling the extent of variation between fibres and along fibres, respectively. The values for these parameters were hand-tuned (values listed in Table 1) to create fibres appropriate for the microfibrotic patterns that we consider here, but can be adjusted (or allowed to vary and tuned using our automated approach) as required for the capture of different patterns.

Conversion from the images 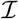 produced by the generator (3) into binary patterns is achieved simply by selecting a threshold value that delineates myocardial tissue and collagenous obstruction. This threshold is selected (via the bisection method) such that the resulting pattern has the desired density of collagen. When matching to experimental image data, the desired density is the density of collagen evident in the target image.

### Automated Generator Tuning

Tuning is achieved by exploring the parameter space of the generator (Table 1) to find patterns that best match the features of the target pattern. Denoting the vector of metrics for a pattern **m** and the target patten **m**\*, we seek to minimise some measure of discrepancy between these metrics, *ρ*(**m**, **m**\*). The generator can produce many different patterns (and therefore, metric values) for a single set of parameters (different noisefield realisations). Therefore, this is a stochastic optimisation problem. One way to express this is to define

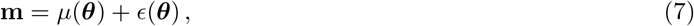

with ***θ*** = (*l*_*b*_, *R*, *γ*, *l*_*d*_, *L*, *ϕ*, *f*, *d*) the set of the generator’s variable parameters, *μ* an unknown function defining the baseline (say, mean) behaviour of the generator function in terms of the metrics, and *ϵ* a random variable whose distribution depends on the parameters.

This thinking permits a Bayesian approach to the problem, but as the structure of *ϵ*(***θ***) is unknown, no likelihood function is available. ABC is a simple and popular likelihood-free technique [24], and here we use the sequential Monte Carlo variation, SMC-ABC, an idea that began with Sisson *et al.* [46]. SMC-ABC attains much higher acceptance rates than traditional ABC, whilst retaining the ease of parallelisation and ability to deal with complex posteriors that are lost by some other advanced ABC variants. A further advantage of ABC in this context is that it explicitly defines the (approximate) posterior in terms of its summary statistics, here the pattern quantification metrics. This way, the posterior samples it generates prioritise matching in terms of the key pattern features.

Without information as to how the generator function behaves and for the sake of simplicity, we choose a uniform prior for all variable parameters. However, as the anisotropy is specified in terms of *R*, which is a ratio, we first transform it logarithmically so that there is equal probability of its value being in the interval [0.5*R, R*] and [*R*, 2*R*]. We use the SMC-ABC algorithm presented by Drovandi and Pettitt [47], with details provided in the supplementary material. Briefly, the algorithm acts by initialising *N* particles across the parameter space, here using Latin hypercube sampling to obtain better coverage [48] at a cost of the initial sample matching the prior only in the marginal sense. Patterns and the corresponding metrics are calculated for each particle, and then the best (in terms of discrepancy from the target pattern) half of the particles are retained. The remaining particles are replaced by copies randomly selected from the retained particles, and then perturbed by Markov chain Monte Carlo (MCMC) steps that seek to find new (and unique) locations for particles that remain within the current limit of allowable discrepancy as defined by the worst kept particle. These two processes of particle copying and perturbation are repeated until the discrepancy shrinks to the desired level, or the perturbation process fails to find sufficient unique locations for particles (suggesting a difficulty to further improve the population). A multivariate normal proposal distribution was used for MCMC steps, with covariance matrix equal to the covariance matrix of the set of particles, scaled by 2.38^2^/*N*_params_ as is common in the literature and originally proposed by Gelman *e*t al. [49].

When the SMC-ABC particles are initialised (as a sample from the prior), the metric values across particles can be used to calculate sample variances *s*_*i*_ for each of the metrics. This enables a weighted Euclidean distance,

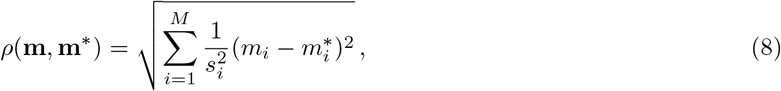

as the discrepancy measure, ensuring that calculated discrepancies are not overwhelmed by individual metrics that naturally vary over large ranges [50]. Mahalanobis distance, which additionally takes into account the correlations between individual metrics [51], was also trialled but was found to occasionally assign inappropriately low discrepancy values to patterns with wholly incorrect orientation, and thus discarded.

We further modify our discrepancy calculations by weighting the differences in orientations of ellipses according to the eccentricity of the corresponding ellipse of the target pattern (see details in Supplementary material), because matching the orientation of a near-circular ellipse is of course far less important than matching the orientation of one that indicates significant anisotropy in the pattern. Differences in orientation also take into account its periodic nature (that is to say, an angle of 89° will be correctly judged as 2° away from an angle of −89°).

### Electrophysiological Simulations

We use computational simulation of the propagation of cardiac excitation to explore how the electrophysiological impact of different microscopic structures can vary both within, and between, the different categories of fibrosis. The ability to consider variability within individual fibrosis categories comes directly from our ability to computationally generate new realisations of a given type of patterning. We simulate under conditions of chronic atrial fibrillation (cAF),

Simulations were carried out in two dimensions, on 4.78mm × 3.68mm (650 × 500 pixels) slices of myocardial tissue. A fibrotic region was added towards the centre of each solution domain, into which the patternings of collagenous obstruction were placed. Pixels occupied by collagen were made non-conductive, a common approach for modelling the impacts of fibrosis on cardiac signal conduction [52, 13]. Fibrotic regions were created via a simple region-growing algorithm that produced regions roughly matching in size with the representative histological sections, but with more naturally-shaped edges (further details in the supplementary material). The impact of the fibrotic region on conduction was considered by stimulating a small area on one side of the tissue, causing signal to propagate both through and around the region of partial obstruction [19, 15]. The simulation setup is also visualised in Figure S5.

Propagation of excitation was modelled using the monodomain formulation [53],

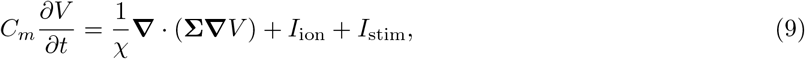

with a conductivity tensor defining a base rate of conduction *σ*_*l*_ along fibres (with orientation *ϕ*) and a 3:1 longitudinal to transverse conductivity ratio,

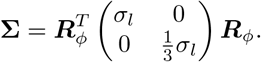

We selected the Maleckar *et al.* cell model [54] for *I*_ion_ as a reasonably modern representation of ion transport in atrial myocytes. Ion channel conductances were modified according to our compilation of experimental data regarding the electrical modelling that takes place under cAF [55, Table 1]. The stimulus current *I*_stim_ was 150,000 *μ*A/cm^−3^, applied for 5 ms at a series of times with reducing cycle length, *t*_stim_ = (200, 550, 900, 1225, 1550, 1850, 2150, 2400, 2650, 2890, 3130, 3360, 3590, 3810, 4030, 4240, 4450, 4650, 4850, 5050, 5250, 5430, 5610, 5770, 5950, 6100, 6250, 6400, 6550, 6700) ms.

The Chaste simulation framework [56] was used for numerical solution of equation (9) via the finite element method, with CVODE [57] used for temporal integration. Node points were placed at the corners of pixels, except where this would cause a node point to be completely submerged within collagenous obstruction. Boundaries were marked along borders between occupied and unoccupied pixels so as to apply no-flux conditions, allowing only unoccupied pixels to be assigned mesh elements. Each unoccupied pixel was divided into two triangular mesh elements (for compatibility with Chaste’s finite element implementation), resulting in meshes of 550,000–650,000 elements. The values of the remaining monodomain parameters, the cell capacitance *C*_*m*_ = 1 *μ*F/cm^2^, surface to volume ratio *χ* = 1400 cm^−1^ and longitudinal conductivity *σ*_*l*_ = 1.75 mS/cm are the Chaste defaults.

## Open Source Code

All MATLAB code used in the production of this paper’s results, including the fibrosis generator and the SMC-ABC algorithm used for parameter tuning, is available on GitHub: https://github.com/betalawson/perlin_microfibrosis

## Author Contributions

D.J., K.B., C.D., A.B.O. and B.A.J.L. contributed to the original development of presented methodology. D.J., C.D. and B.A.J.L. contributed to the computer implementation of methods. K.B., P.B., A.B.O., R.W.S., B.R. and B.A.J.L. contributed to the analysis of generated patterns and electrophysiological simulations. All authors contributed to the drafting and refinement of the manuscript.

## Acknowledgements and Funding

K.B., P.B. and B.A.J.L acknowledge financial support provided by the ARC Centre of Excellence for Mathematical and Statistical Frontiers (CE/140100049). A.B.O. acknowledges a British Heart Foundation Intermediate Basic Science Fellowship (FS/17/22/32644). R. W. S. acknowledges the support of Universidade Federal de Juiz de Fora (JFJF), and funding bodies Conselho Nacional de Desenvolvimento Científico e Technológico, Co-ordenação de Aperfeicoamento de Pessoal de Nível Superior and Fundaçãao de Amparo à Pesquisa do Estado de Minas Gerais. B.R. acknowledges a Wellcome Trust Fellowship in Basic Biomedical Sciences (100246/Z/12/Z and 214290/Z/18/Z). A.B.O. and B.R. also acknowledge funding from an Impact for Infrastructure Award from the National Centre for the Replacement, Reinfement and Reduction of Animals in Research (NC/P001076/1). The authors would also like to acknowledge Queensland University of Technology’s eResearch Office, for their assistance with the installation of Chaste on high power computing architecture and the use of these facilities to perform electrophysiological simulations.

