## Supplementary Material for "Perlin Noise Generation of Physiologically Realistic Patterns of Fibrosis"

### Additional Information/Methods

#### Perlin Noise and its Implementation

Perlin noise is a type of *gradient noise*, so called because it is constructed using a series of gradient vectors, specifically randomly-oriented unit vectors assigned with each point in a grid of regular, unit spacing. Important properties of the technique include the very natural extension to any number of dimensions, and the potential for implementations that scale as $O(N_{\text{points}})$ by approximating the random property of the gradient vectors. We describe the specifics of our implementation after first describing the method in detail, presenting the two-dimensional formulation of Perlin noise used in this work.

The value of a Perlin noise field $\mathcal{P}$ at a given point in space, $\mathbf{x}$, is given by the smoothly interpolated combination of a set of dot products between pairs of vectors. The first vector in each pair is the vector connecting $\mathbf{x}$ to one of its surrounding gridpoints $\mathbf{x_i}$, denoted here $\mathbf{r}_i = \mathbf{x} - \mathbf{x}_i$. The second vector in each pair is the gradient vector associated with that gridpoint, denoted here $\mathbf{g}_i$. In two dimensions, there are four surrounding gridpoints, and the noisefield takes on a value

$$\mathcal{P}(\mathbf{x}) = \sum_{i=1}^{4} \alpha\big(1 - |(\mathbf{r}_i)_x|\big)\alpha\big(1 - |(\mathbf{r}_i)_y|\big)(\mathbf{r}_i \cdot \mathbf{g}_i), \tag{10}$$

with $(\mathbf{r}_i)_x$ denoting the $x$ co-ordinate of $\mathbf{r}_i$ and $\alpha(x)$ a scalar function that controls the smoothing. We use here the improved form suggested by Perlin [58],

$$\alpha(x) = 6x^5 - 15x^4 + 10x^3.$$

Due to the unit grid, the inputs of interest for the smoothing function are $x \in [0, 1]$, and over this domain its range is also $[0, 1]$. The function has zero derivatives at both ends of its domain, and is symmetric in the sense that it satisfies the property $f(1-x) = 1 - f(x)$. This property is what permits the concise expression of Perlin noise given in equation (10), but the process is better understood as a linear interpolation applied first in one dimension, and then the other, with $\alpha$ a smoothing transformation applied to each co-ordinate in turn. $\alpha(x) = x$ thus recovers traditional (bi)linear interpolation. In-depth descriptions of Perlin noise and the use of this smoothing function are readily available online.

As is typical in practise, our implementation of Perlin noise does not generate random vectors $\mathbf{g}_i$ for each gridpoint. Instead, 256 vectors equally spaced around the unit circle are pre-generated, as well as permutation tables containing a random permutation of the numbers 0 to 255 for each octave of Perlin noise. Vectors are then pseudo-randomly selected from the pre-generated set as required via a hashing algorithm, using the permutation tables to essentially randomise the initial inputs. Specifically, using the bottom-left grid corner as an example, a vector is selected by flooring the $x$ and $y$ co-ordinates of the evaluation point, then finding their value modulo 256. One of these two values is fed into the permutation table to randomise it, then the result is hashed with the other via a bitwise XOR operation to generate a new value in the range 0 to 255 that can be fed into the permutation table. Vectors for the other three corners are found the same way, except with a ceiling operation instead of a floor operation as appropriate. For Perlin noise as used in this work, the random offsets between different octaves are also pre-generated. When a new realisation of Perlin noise is desired, it can be created simply by generating new permutation and offset tables, re-randomising the gridpoint vectors and thus creating a new noisefield but with the same underlying properties.

#### Noisefield Example Images

Figure 7 illustrates different noise fields, and some of the concepts we use to obtain different types of patterning. The methods used to generate these points are presented in Table S1.

#### Sequential Monte Carlo Approximate Bayesian Computation

Approximate Bayesian computation (ABC) is a technique for sampling from an approximation to a posterior distribution implied by a set of data and imposed likelihood structure. The method is particularly important in cases where the likelihood is very expensive to calculate, or indeed computationally intractable as is the case here. The method works by generating samples, then rejecting those that fall too far from the data. Typically the data is high-dimensional (here the pixels in patterns), and so discrepancy from the target data is considered in terms of a much smaller set of summary statistics (here the pattern metrics). In the traditional form of ABC, samples are simply generated from the prior, then accepted or rejected according to falling within a pre-specified level of acceptable discrepancy.

| Noisefield | Description |
|---|---|
| White Noise | Uniform value on [0,1] at every pixel |
| Single-octave Perlin Noise | One octave of Perlin noise with feature size $l = 48$ pixels |
| Multiple-octave Perlin Noise | Perlin noise with base feature size $l = 48$ pixels, four octaves with weighting $\gamma = 0.5$ |
| Anisotropic Perlin Noise | As in (iii), but with a rotation of $\phi = \pi/3$ and squeeze mapping with $R = 3.5$ applied, see equation (2) in main document |
| Additive Combination | As in (iv), but combined additively with a second isotropic Perlin noise field with $l = 400$ pixels, four octaves with weighting $\gamma = 0.5$ (60% original, 40% second field) |
| Gaussian Random Field | Gaussian random field generated using circulant embedding [59], with a Matern-3/2 covariance function and $l = 15$ pixels. |

Table S1: The different noisefields visualised in Figure 7.

Sequential Monte Carlo (SMC) is a family of sampling techniques that uses a population of particles that move through the parameter space, acting as samples from a series of intermediary distributions that approach the posterior distribution from which samples are desired. In this manner, the complexity of the sampling problem is gradually introduced. The ABC version of SMC forms the intermediary distributions by changing the level of discrepancy required for sample acceptance. By choosing these discrepancy levels in response to the current population of particles, complexity is introduced in a manner that ensures it will not be too difficult to generate acceptable samples, except when sample quality is no longer improving and the method can be terminated. This dynamic determination of when to stop the method also means the acceptable level of discrepancy need not be pre-specified, the method instead generating samples of lower and lower discrepancy until it appears unable to do better.

The method is initialised by placing a pre-specified number of particles, at locations in the parameter space sampled from the prior. Data associated with each particle is generated, here by running the generator using those values for its parameters. Metrics are calculated for each of these patterns, and then an acceptable level of discrepancy for the next sampling step is selected using the median discrepancy (worst discrepancy among the best half of particles). In order to generate samples that are likely to satisfy this discrepancy requirement, the particles that do not already satisfy it are copied (randomly with replacement) onto the particles that do. New samples are then generated by attempting to move the copied particles to nearby locations, and only accepting these moves when the generated pattern associated with the new location has a satisfactory discrepancy from the target. After a dynamically chosen number of move steps [47], the particles (potentially with some surviving duplicates) are now a sample from the new distribution. The process then repeats, each step updating the acceptable discrepancy using the median particle and copying and moving the particles that do not satisfy the new discrepancy value, until the move steps do not successfully generate a sufficient number of unique particles. The method is laid out in detail in Algorithm 1.

**Metric Weighting**

The metrics used to judge discrepancy between different patterns are the orientations and axis lengths (major and minor) of a set of ellipses, obtained from the smoothed power spectrum corresponding to a given pattern. As discussed in the main document, an initial sample allows for the variance of each of these metrics to be estimated, and thus they can be scaled so that each varies over a similar region and none is able to bias calculated discrepancy values. However, a further correction improves calculated discrepancies, namely in the case where an ellipse(s) is approximately circular. In this case, the orientation is essentially meaningless, and in general a failure to match an ellipse in orientation should be considered more severe when that orientation carries with it a significant anisotropy. For this reason, when calculating the discrepancy from some target we weight differences in orientation $\phi$ by an additional factor,

$$\Delta\phi = \ln\left(\frac{a}{b}\right)\Delta\phi_{\text{orig}},$$

where $a$ and $b$ are the lengths of the major and minor axes of the corresponding ellipse in the target, respectively. This approach results in an unmodified value when the anisotropy ratio of the relevant ellipse is $e \approx 2.71828$, upweighting or downweighting the contribution when the anisotropy is stronger or weaker than this value, respectively. Most importantly, this modification correctly results in a contribution of zero to discrepancy from an ellipse's orientation when the target ellipse is a circle ($a = b$) and its orientation is therefore arbitrary.

**Algorithm 1** SMC-ABC algorithm

▷ Define number of particles to keep for each step
$K \leftarrow \text{round}(\alpha N)$ ▷ Here $\alpha = 0.75$

▷ Initialise particles
Sample particle locations, $\boldsymbol{\theta}$, via Latin hypercube sampling
Generate patterns associated with all particles, $\boldsymbol{\Gamma}$
Calculate weightings for each metric in discrepancy calculations (Eq. 8 in main document)
Calculate discrepancies $\epsilon$ between each pattern $\boldsymbol{\Gamma}$ and the target pattern $\boldsymbol{\Gamma}_{\text{target}}$

▷ Loop until particle improvement grows too slow
**while** !`stop` **do**

▷ Select target discrepancy for this step according to worst kept particle
Sort particles according to descending $\epsilon$ value
$\epsilon_{\text{target}} \leftarrow \epsilon_\alpha$

▷ Copy unsatisfactory particles onto satisfactory particles
**for** $i \leftarrow K+1$ to $N$ **do**
$r \leftarrow$ random integer in $[1, K]$
$\boldsymbol{\theta}_i \leftarrow \boldsymbol{\theta}_r$
$\boldsymbol{\Gamma}_r \leftarrow \boldsymbol{\Gamma}_i$
**end for**

▷ Perform single move step on copied particles
`success` $\leftarrow 0$
**for** $i \leftarrow K+1$ to $N$ **do**
▷ Trial move
Sample $\boldsymbol{\theta}' \sim N\left(\boldsymbol{\theta}_i, \frac{2.38^2}{N_{\text{params}}}\text{Covar}(\boldsymbol{\theta})\right)$
Generate pattern, $\boldsymbol{\Gamma}'$, using $\boldsymbol{\theta}'$ and calculate its discrepancy, $\epsilon'$
▷ Accept if discrepancy is satisfactory
**if** $\epsilon' \leq \epsilon_{\text{target}}$ **then**
$\boldsymbol{\theta}_i \leftarrow \boldsymbol{\theta}', \ \boldsymbol{\Gamma}_i \leftarrow \boldsymbol{\Gamma}', \ \epsilon_i \leftarrow \epsilon'$
`success` $\leftarrow$ `success` $+1$
**end if**
**end for**
▷ Calculate proportion of successful moves
Set $s \leftarrow$ `success`$/(N-K)$

▷ Perform the remaining of a dynamically selected number of moves
$R \leftarrow \max\left(\text{ceil}\left(\frac{\ln 0.05}{\ln(1-s)}\right), 300\right)$
**for** $j \leftarrow 1$ to $R-1$ **do**
**for** $i \leftarrow K+1$ to $N$ **do**
Sample $\boldsymbol{\theta}' \sim N(\boldsymbol{\theta}_i, \frac{2.38^2}{N_{\text{params}}}\boldsymbol{\Sigma})$
Generate pattern, $\boldsymbol{\Gamma}'$, using $\boldsymbol{\theta}'$ and calculate its discrepancy, $\epsilon'$
**if** $\epsilon' \leq \epsilon_{\text{target}}$ **then**
$\boldsymbol{\theta}_i \leftarrow \boldsymbol{\theta}', \ \boldsymbol{\Gamma}_i \leftarrow \boldsymbol{\Gamma}', \ \epsilon_i \leftarrow \epsilon'$
**end if**
**end for**
**end for**

▷ Terminate when particle degeneracy implies a lack of further improvement
**if** $N_{\text{unique}} < N/2$ **then**
`stop` $\leftarrow$ `true`
**end if**

**end while**

#### Region Growing Algorithm

The generated patterns can be directly used in cardiac electrophysiology simulations, however the rectangular domain (chosen to match the histological images they are based on) may result in unrealistically sharp edges at the boundaries of the fibrotic region. In order to create more naturally-shaped regions, the rectangular domains on which the histological and generated patterns are defined were first reduced to softer-edged (but still approximately rectangular) shapes via a simple region growing algorithm. This algorithm creates regions like the one marked in grey in Figure S5, into which the actual pattern of fibrotic obstruction is then inserted (dark red colour in figure).

The algorithm initialises with a set of “seeds”, which are locations that are marked as belonging to the reduced shape. The positions of these seeds in our use of this algorithm are shown in Figure S5. By placing these seeds towards the centre of the domain, it grows outwards from each, almost surely filling in the areas between seeds but resulting in a rougher and less rectangular region boundary. Growth of the region proceeds by repeatedly selecting a random grid location that has already been marked as part of the region (on the first step, this will be one of the seeds), then picking one of its neighbours (von Neumann neighbourhood) at random. If that neighbour is not yet part of the region, it becomes part of the region. This process then repeats until a specified number of sites are included in the region, here 85% of the original rectangular domain.

### Supplementary Figures

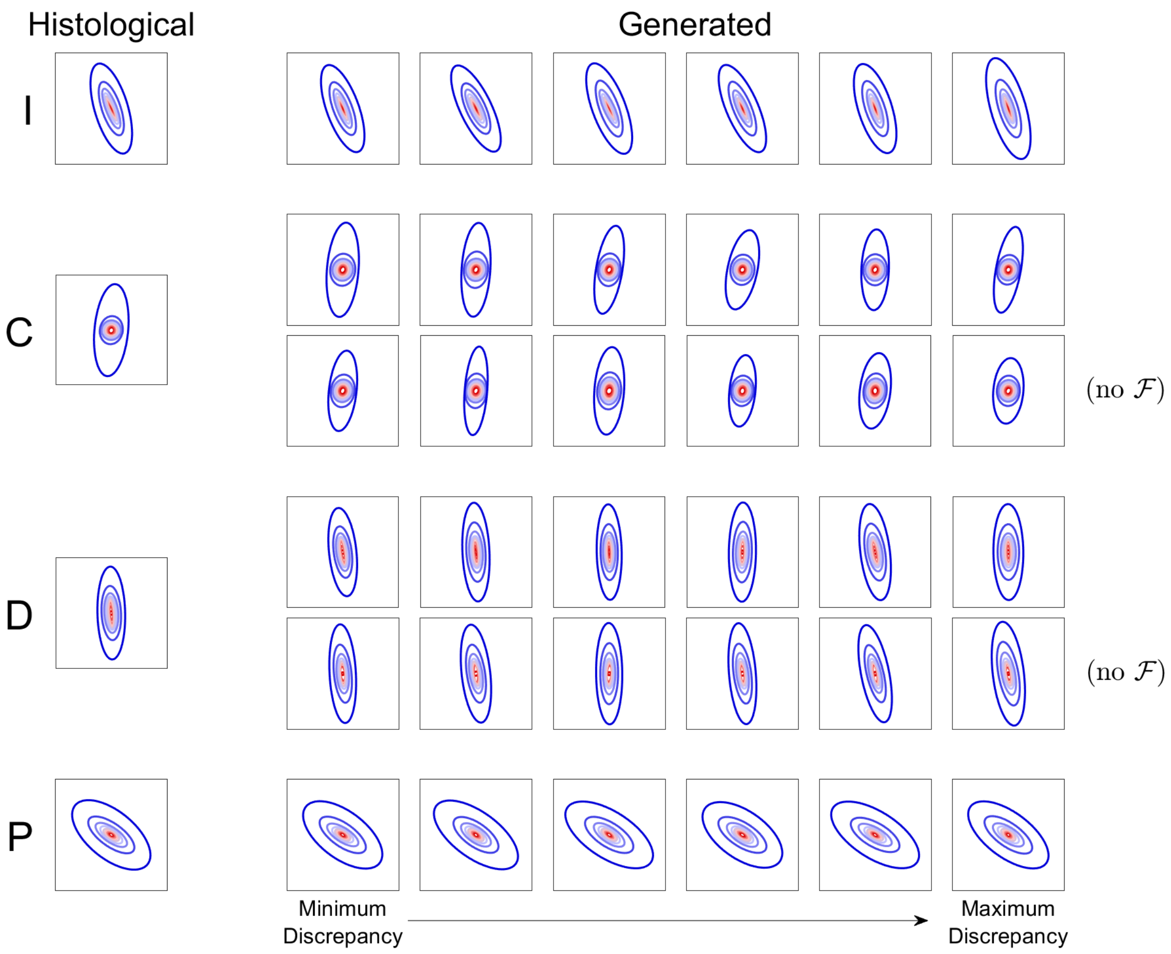


Figure S1: Successful establishment of low-discrepancy populations matched to the histological patterns. Metric ellipses corresponding to the patterns visualised in Figure 4 of the main document, which are the patterns matched to the different types of cardiac fibrosis, interstitial (I), compact (C), diffuse (D) and patchy (P). Axes are consistent within (but not across) the different classifications of microfibrosis. In all cases, the metric ellipses of the generated patterns match very well with the histological target, including the cases where the fibre-selecting field was not included in the automatic tuning (no $\mathcal{F}$). This indicates the success of the SMC-ABC algorithm in generating low-discrepancy patterns.

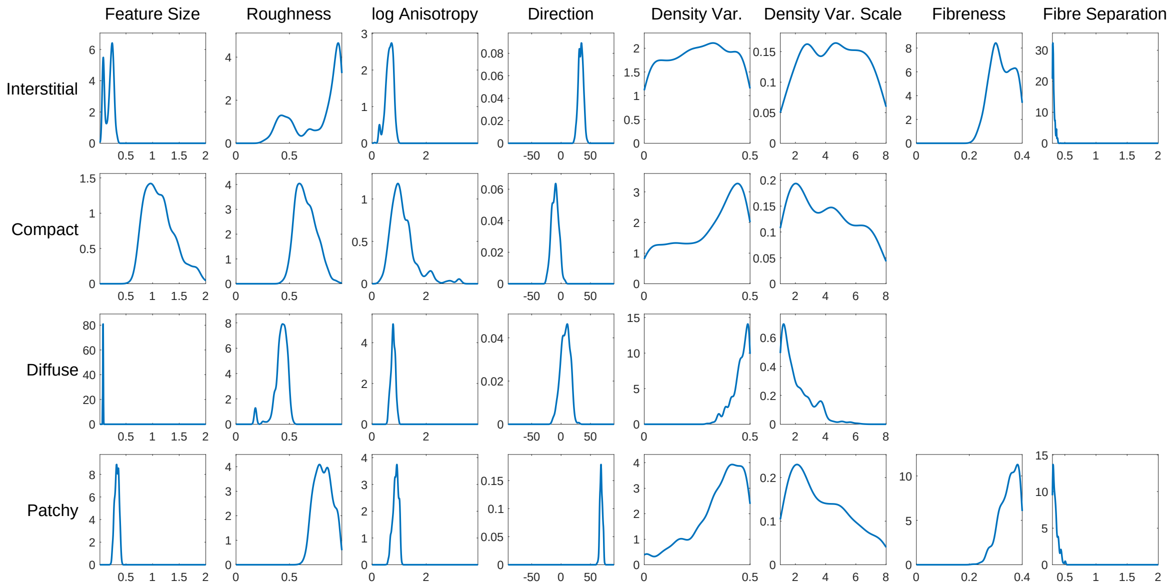


Figure S2: Generator parameters reproducing microfibrosis structure. Marginal distributions of the generator parameter values in the populations constructed to represent the four different classes of fibrosis. Although some distributions are complex, the automatic selection of parameter values by the SMC-ABC process largely results in what could be expected by considering tuning parameter functions in the context of the patterns being matched. For example, the smallest features are selected for the diffuse patterns, then interstitial and patchy, and finally the largest features are selected for compact fibrosis. Further detail is provided in the text.

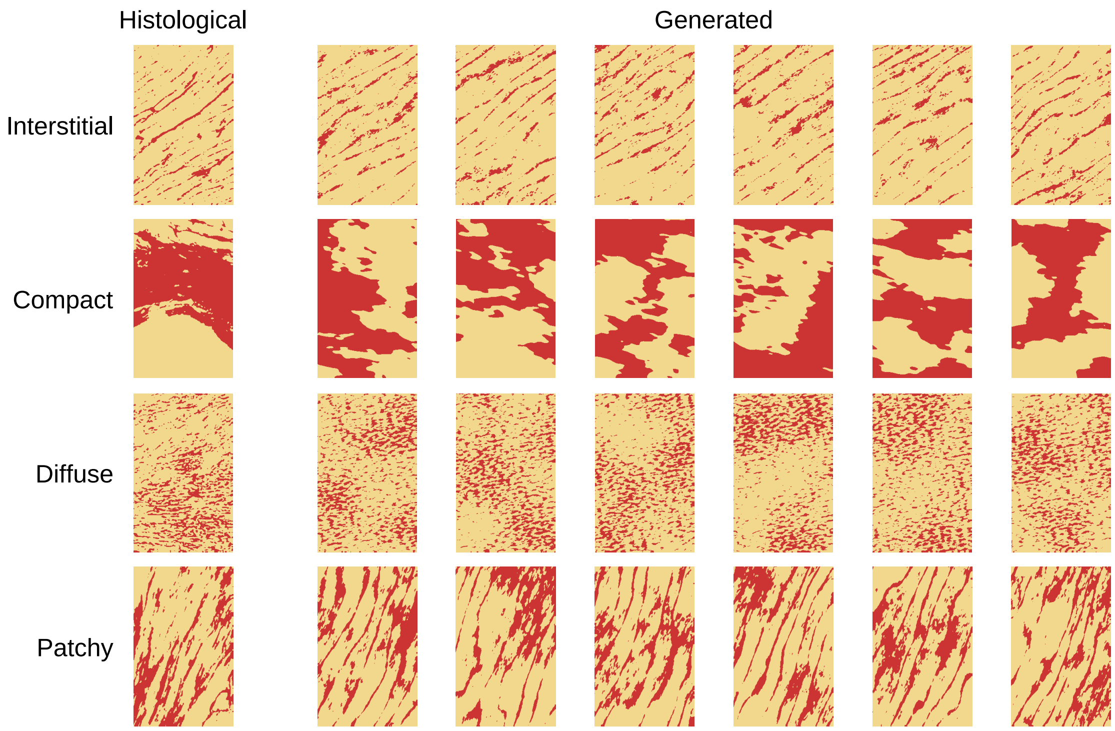


Figure S3: Patterns generated using the proposed generator tunings. Sample patterns generated by running the generator with parameter values as defined in Table 2, and random seed information. Visual agreement with the histological sections is consistently demonstrated, verifying the use of these singular parameter sets in generating further realisations of the different kinds of microfibrotic patterning.

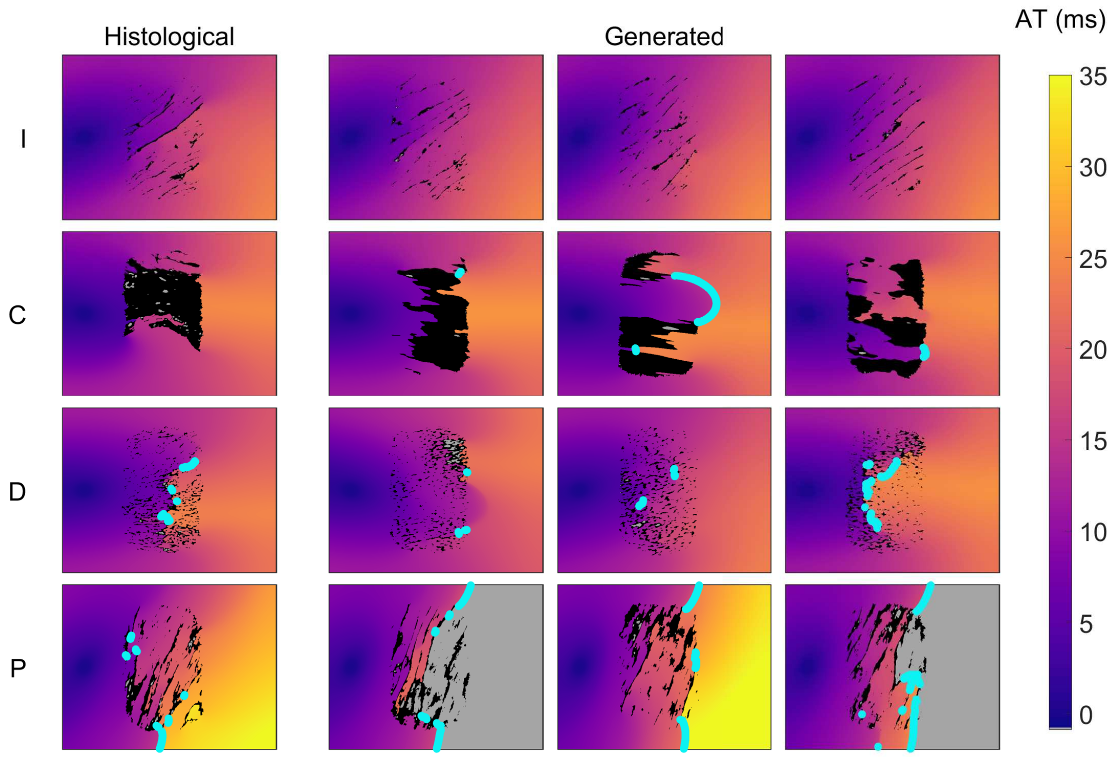


Figure S4: Activation maps arising due to fibrotic regions with different microfibrotic patterning. The activation times visualised are those corresponding to the same stimuli shown in Figure 5 in the main document. Lines of significant conduction block are now marked in bright blue.

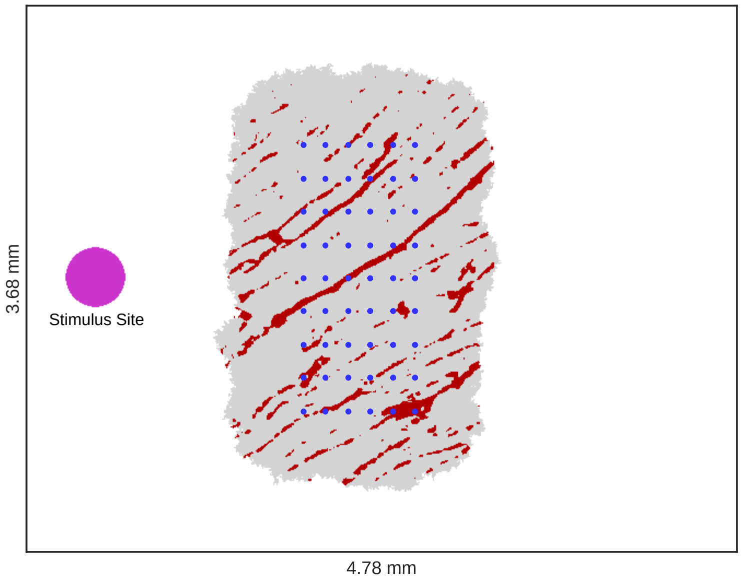


Figure S5: The simulation setup. Schematic diagram of the simulation domain, showing how a region of fibrosis with defined microfibrotic structure is incorporated. The grey area shows the region generated by the region growing algorithm, in which the pattern of collagenous obstruction (dark red) is placed. Blue dots indicate the placement of the "seeds" used by the region growing algorithm (see Supplementary material). Pixels not occupied by collagen have the same properties, regardless of falling outside (white) or inside (grey) the region marked as fibrotic.
